# Leveraging ONT move table values for signal aware variant calling

**DOI:** 10.64898/2026.02.13.705285

**Authors:** Xian Yu, Zhenxian Zheng, Lei Chen, Zilan Qin, Minggao He, Ruibang Luo

**Affiliations:** School of Computing and Data Science, University of Hong Kong, Hong Kong, China

**Keywords:** Oxford Nanopore sequencing, small variant calling, move table

## Abstract

Oxford Nanopore Technologies (ONT) sequencing enables long-range haplotype phasing and contiguous genome assembly but still exhibits elevated error rates that challenge small variant calling, particularly for insertions and deletions (Indels). While raw electrical signals contain rich information, existing signal-aware methods require computationally intensive processing of large signal files. Here, we present Clair3 v2, a method that leverages the ONT move table—a lightweight byproduct of basecalling that maps signal events to nucleotide positions—to improve variant calling accuracy. Clair3 v2 builds upon Clair3 and integrates signal-level dwelling time to significantly enhance variant calling performance. We also propose a genome position based circular buffer to incorporate dwelling time with minimal computational overhead. Benchmarking across six Genome in a Bottle samples demonstrates substantial improvements in variant calling accuracy. With HAC basecalling, Clair3 v2 achieves a mean SNP F1-score of 97.69% at 10× depth (compared to 96.45% for baseline Clair3), and Indel F1 scores improved from 64.27% to 76.70%, while gains persisted at higher depths. The benefits were most pronounced for longer Indels and in complex genomic regions, where Indel F1 scores in long homopolymer regions improved from 14.3% to 45.2%. Benchmark results across various basecalling modes, samples, and coverage settings outperformed Clair3 baselines and other methods, including DeepVariant and Dorado Variant, and demonstrate the significant benefits of Clair3 v2. Furthermore, Clair3 v2 incurs negligible runtime compared to standard Clair3, making it practical for routine use.

## 1. Introduction

Oxford Nanopore Technologies (ONT) has emerged as one of the mainstream long-read sequencing platforms. By measuring electrical current signals as DNA molecules pass through a nanopore, ONT enables real-time, portable long-read sequencing that supports diverse downstream applications, including direct detection of DNA modifications (Ahsan et al., 2024), structural variant detection (Smolka et al., 2024, Jiang et al., 2025), and improved genome assembly (Chen et al. 2021, Cheng et al., 2026). Recent advances in sequencing chemistry (e.g., R10.4.1) and basecalling models have substantially improved accuracy (Sereika et al., 2022), narrowing the gap with short-read technologies. Despite these improvements, accurate detection of insertions and deletions (indels) remains a significant challenge, particularly in low-complexity genomic regions.

State-of-the-art variant calling tools such as Clair3 (Zheng et al., 2022), DeepVariant (Poplin et al., 2018), and LongcallD (Gao, 2025) achieve competitive SNP performance but struggle with indels, especially in repetitive sequences (Goenka et al., 2022). These methods rely on alignment features, leaving raw signal information untapped. The ONT signal encodes the physical duration of DNA translocation through the pore, providing kinetic information that is particularly valuable for distinguishing true variants from artifacts. The ONT move table, a lightweight byproduct of basecalling that maps signal events to basecalled positions, remains underexplored for variant calling despite its proven utility for modification detection (Luo et al., 2025). While Dorado (Oxford Nanopore Technologies, 2024) leverages move-table data for polishing and variant calling, its whole-genome polishing approach is computationally intensive, motivating the need for efficient, signal-aware variant calling methods.

In this work, we present Clair3 v2, an enhanced version of Clair3 that incorporates move-table–derived dwelling times as additional features in the full-alignment model. Benchmarking across six Genome in a Bottle (GIAB) samples demonstrates that incorporating dwelling time substantially improves variant-calling accuracy across all tested sequencing depths (10× –50×). For SNPs, mean F1 scores improved from 96.45% to 97.69% at 10× and from 99.62% to 99.77% at 50× coverage. The improvements were more pronounced for Indels, with mean F1 scores increasing from 64.27% to 76.70% at 10× and from 83.66% to 89.51% at 50× coverage. Clair3 v2 also outperformed state-of-the-art variant callers, including DeepVariant, Dorado Variant, and LongcallD, achieving comparable or higher F1 scores across all tested conditions while maintaining comparable runtime to the standard Clair3 workflow by using a genome position based circular buffer.

## 2. Methods

### 2.1. Overview of signal-aware Clair3

Our method builds on Clair3, a mature and officially supported ONT variant-calling pipeline that combines an efficient pileup model with a highly accurate full-alignment model (Figure 1). The pipeline takes an aligned BAM file and the corresponding reference genome as input. First, pileup features are constructed and fed into the pileup model to generate an initial set of variant calls. High-confidence variants are then used for read phasing and haplotagging with WhatsHap (Martin et al., 2016) or LongPhase (Lin et al., 2022). For sites with low-confidence calls, together with additional candidate sites selected by heuristic filtering, full-alignment tensors are constructed and passed to the full-alignment model for prediction. Finally, outputs from both models are merged to produce the final variant call set.

**Figure 1.**
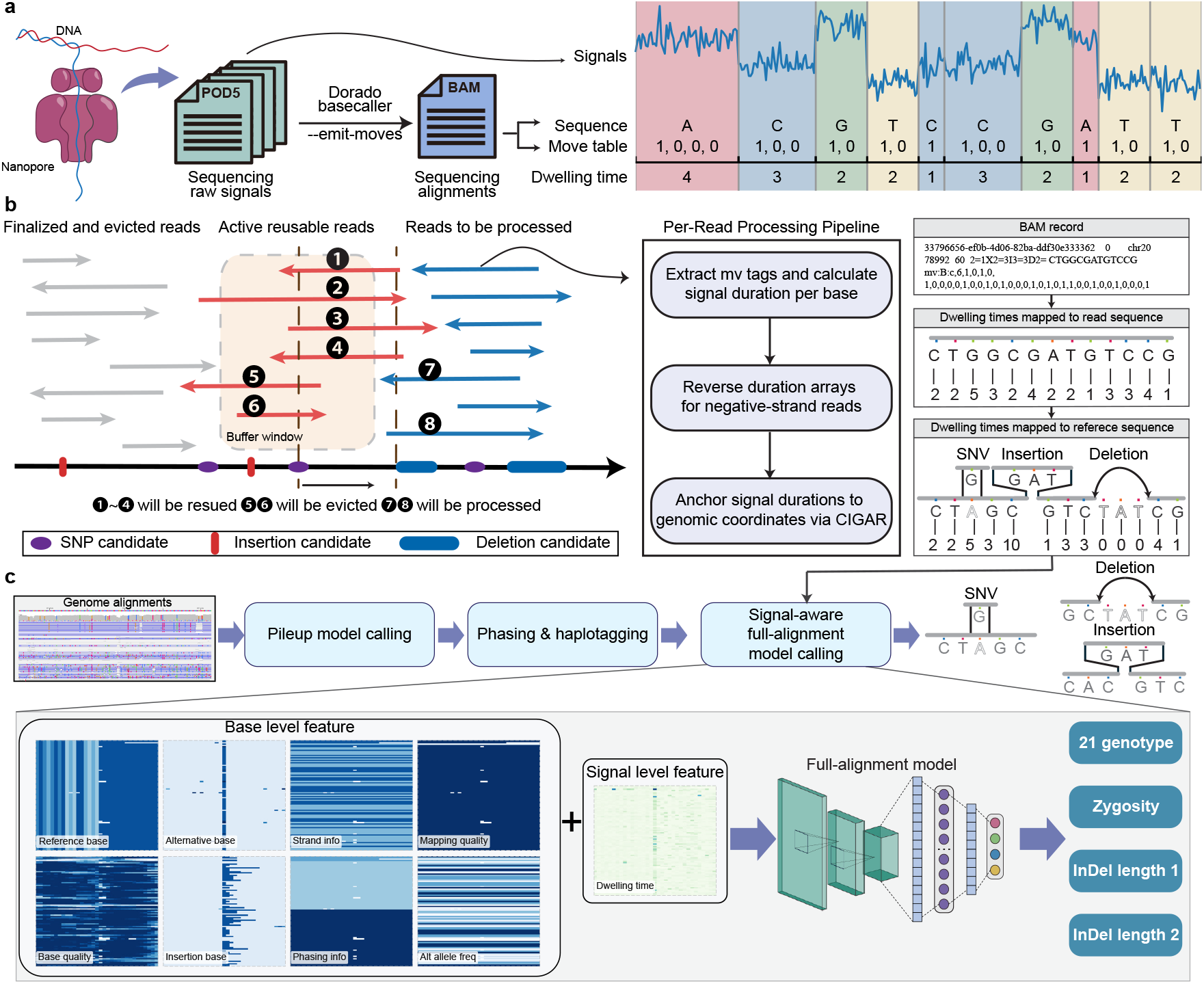
Overview of signal-aware variant calling. **(a)** Dwelling time extraction from move table. Dorado basecaller with `--emit-moves` produces BAM files containing binary signal-to-base mappings, converted to per-base dwelling times. **(b)** Genome position based circular buffer maintains active reads for efficient feature construction. **(c)** Clair3 v2 pipeline integrating base-level and dwelling time features for variant prediction.

Clair3 v2 extends this framework by integrating signal-level dwelling time as an additional feature. Combined with a genome position based circular buffer for efficient feature construction, this approach improves SNP and indel detection accuracy with negligible additional computational cost.

### 2.2. Extracting dwelling time and integrating signal features into full-alignment tensors

Our method leverages the move table generated by the Dorado basecaller when invoked with the `--emit-moves` argument (Figure 1a). During basecalling, raw electrical signals from POD5 files are segmented into discrete events, and the move table records which events correspond to base transitions. The resulting BAM file stores the move table under the `mv:B:c tag`, encoding a stride value *s* followed by a unary sequence **m** = (*m*_1_, *m*_2_, …, *m*_*n*_), where *m*_*k*_ ∈ {0, 1}. The stride indicates the number of raw signal samples per move-table entry (typically 5 or 6 depending on the basecalling model). Each 1 marks a base transition, while 0s indicate that the signal dwelled on the current base. For basecalled nucleotide *i*, its dwelling time *d*_*i*_ is computed as:

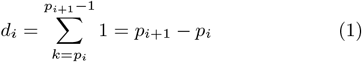

where *p*_*i*_ denotes the position of the *i*-th base transition (i.e., the *i*-th occurrence of *m*_*k*_ = 1) in the move-table sequence. The corresponding raw signal duration is *d*_*i*_ × *s* samples. For example, a move-table entry `mv:B:c,6,1,0,1,0` with stride *s* = 6 indicates that the first base has *d*_1_ = 2 (spanning positions 1–2), corresponding to 12 raw signal samples. We convert these unary sequences into integer dwelling times by counting consecutive entries per base. This approach extracts signal-level information directly from the BAM file without requiring access to the original POD5 files, which are often hundreds of gigabytes in size and slow to access randomly.

Per-read processing proceeds in three steps. First, we extract the `mv tag` from each BAM record and compute the dwelling time for each basecalled nucleotide by summing move-table entries between consecutive base transitions. Second, for reverse-strand reads (indicated by the SAM flag), we reverse the dwelling-time array to match the aligned read orientation, as the move table is stored in the original sequencing direction regardless of alignment. Third, we anchor the per-base dwelling times to reference-genome coordinates using the CIGAR string.

When mapping dwelling times onto the reference, different alignment operations are handled distinctly to create informative patterns for variant calling. Let *D*_*j*_ denote the dwelling time feature assigned to reference position *j*, and *d*_*i*_ the dwelling time of read base *i* aligned to that position:

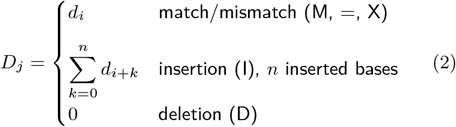

For matches and mismatches, we directly assign the dwelling time of the corresponding read base. For insertions, we sum the dwelling times of the anchor base (*d*_*i*_) and all *n* inserted bases (*d*_*i*+1_, …, *d*_*i*+*n*_), producing elevated values that signal additional bases. For deletions, we assign 0 at each deleted reference position, creating a distinctive pattern for the missing base. For SNPs, dwelling times remain similar to neighboring positions since only the base identity changes. These distinct signatures—elevated for insertions, zero for deletions, and unchanged for SNPs—provide informative features for variant calling.

To integrate dwelling time into the variant calling pipeline, we extend the Clair3 full-alignment tensor by appending dwelling time as an additional channel, expanding its dimension from *L × D × C* (flanking window size *×* read depth *×* feature channels) to *L × D ×* (*C* + 1). At each position, dwelling time values computed according to Equation 2 are populated into this channel. The augmented tensor is then passed to the ResNet-based full-alignment model for variant prediction.

### 2.3. Reducing redundant computation via genome position based circular buffer

In the feature generation stage, candidate sites are evaluated across multiple reads at each genomic position. However, computing dwelling-time features requires parsing the move table at the read level, which would be prohibitively expensive if repeated independently for each candidate position. To bridge this mismatch between position-level candidate detection and read-level feature extraction, we propose a genome position based circular buffer that enables candidate detection and full-alignment feature construction in a single streaming pass over the genome (Figure 1b).

As the scanner advances through genomic coordinates, the circular buffer maintains three categories of reads. *Active reusable reads* have already been parsed and overlap the current position; these are retained for reuse at nearby candidates. *Reads to be processed* are newly encountered reads overlapping the current position that have not yet been parsed; they are processed and added to the active buffer. *Finalized and evicted reads* have alignment end coordinates before the current scanner position and can no longer contribute to future candidates; they are removed to free memory.

During operation, the buffer iterates through genomic positions in sorted order, identifying candidate sites based on preset allele frequency and depth thresholds. When a position is flagged as a candidate, the full-alignment tensor is constructed by gathering features from all covering reads. Pre-computed dwelling-time features are retrieved directly for reads already in the buffer, while newly encountered reads undergo the three-step processing (move-table extraction, strand-aware reversal, and CIGAR-based anchoring) and are cached. As illustrated in Figure 1b, when the genome position based circular buffer advances from one candidate to the next, reads 1–4 that still overlap are reused, reads 5–6 that no longer overlap are evicted, and reads 7–8 that newly overlap are processed and buffered.

This design achieves an efficient runtime-memory trade-off: each read is parsed exactly once regardless of how many candidate positions it spans, while the memory footprint remains bounded by the product of maximum read length and local coverage depth. In practice, this approach adds negligible overhead compared to standard Clair3, as demonstrated in our computational benchmarks.

### 2.4. Model training and performance evaluation

We employed two training configurations. For the dwelling time ablation study, models were trained on GIAB HG002 (chromosomes 1–22, excluding chromosome 20), with the remaining samples (HG001, HG003–HG007) reserved for benchmarking. For the release version, the training set was expanded to include HG001, HG002, and HG005, with chromosome 20 withheld for cross-variant-caller evaluation.

Training data included alignments to both GRCh38 and CHM13 (Hansen et al., 2025) references. Sequencing data were downsampled to 10×, 20×, 30×, 40×, 50×, and 60× coverage using mosdepth (Pedersen and Quinlan, 2018) and samtools (Li et al., 2009). Truth variants were normalized with Repun (Zheng et al., 2025) to ensure consistent variant representation. All data were basecalled using Dorado v5.2.0 with HAC and SUP models. Two methodological notes warrant mention. First, although HG002, HG003, and HG004 belong to the same family trio, consistent improvements on unrelated samples (HG005–HG007) suggest that our method captures generalizable signal patterns rather than sample-specific features (see **Results** section). Second, the Clair3 models in this study differ from ONT’s officially released models, which use different training data and configurations. ONT-provided models typically outperform our baseline models, as our training here is limited to HG002. Detailed data generation procedures are provided in the Supplementary Materials.

The pileup model uses a bidirectional LSTM to predict genotype classes (21 possible allele combinations across two haplotypes) and zygosity states (homozygous, heterozygous, or reference). The full-alignment model employs a ResNet architecture and additionally predicts indel length. For full-alignment training, BAM files were phased with WhatsHap or LongPhase and haplotagged using truth variants. Input tensors incorporated phasing information and dwelling-time features for candidate sites identified by allele frequency and depth thresholds. Both models were trained with Focal Loss and Rectified Adam. Hyperparameter settings are detailed in the Supplementary Materials.

Variant calling performance was evaluated using hap.py, qfy.py (Eberle et al., 2017), and RTG Tools vcfeval (Cleary et al., 2015) against GIAB high-confidence truth sets on GRCh38. hap.py and vcfeval provided haplotype-aware variant comparison, while qfy.py generated stratified performance metrics across genomic regions. Precision, recall, and F1-score were computed separately for SNPs and indels.

## 3. Results

### 3.1. Performance in various basecalling and coverage settings

For fair comparison, we trained two models—Clair3 v2 and baseline Clair3—using identical training data (HG002 sample, excluding chromosome 20) and benchmarked both across six independent samples (HG001, HG003–HG007) at sequencing depths ranging from 10× to 50× (Figure 2).

**Figure 2.**
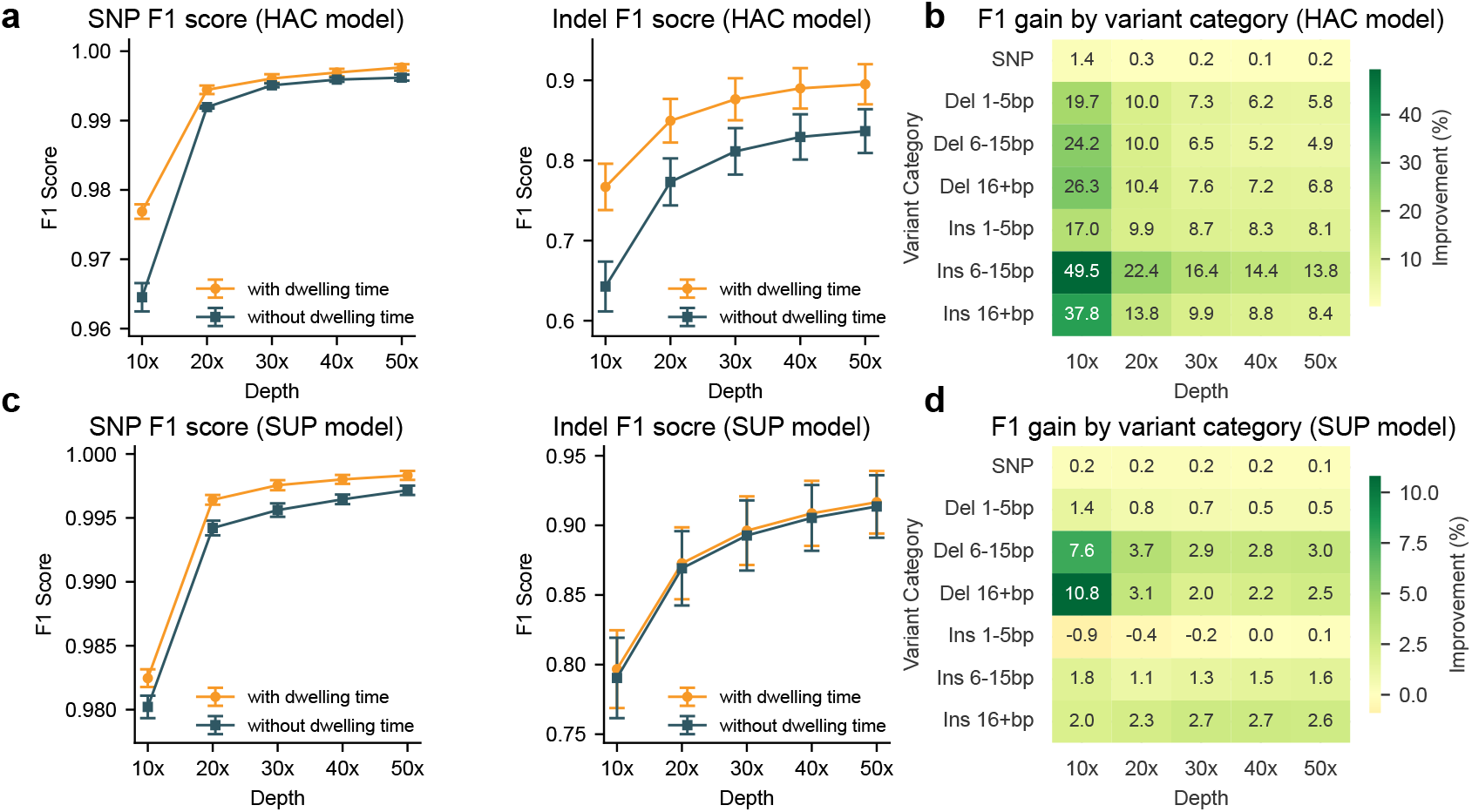
Variant calling performance across sequencing depths. **(a, c)** SNP and Indel F1 scores for HAC and SUP basecalled reads, respectively. Error bars: s.d. across six GIAB samples. **(b, d)** F1 improvement heatmaps by variant category (SNPs, deletions, insertions stratified by length) for HAC and SUP reads on HG005.

Clair3 v2 consistently improved variant calling accuracy across all conditions, with the largest gains observed at lower coverage (Figure 2a,c; Supplementary Tables 1–2). For HAC basecalled reads at 10× depth, mean indel F1 scores improved from 64.27% to 76.70% (s.d. 3.11% versus 2.89%; n = 6 samples), and mean SNP F1 scores increased from 96.45% to 97.69%. These improvements persisted at 50× depth, where indel F1 scores rose from 83.66% to 89.51% and SNP F1 scores from 99.62% to 99.77%. SUP basecalled reads, which have higher baseline accuracy, showed smaller but consistent improvements (Figure 2b,d; Supplementary Tables 3–4): indel F1 scores improved from 79.04% to 79.67% at 10× and from 91.35% to 91.66% at 50×.

**Table 1.**
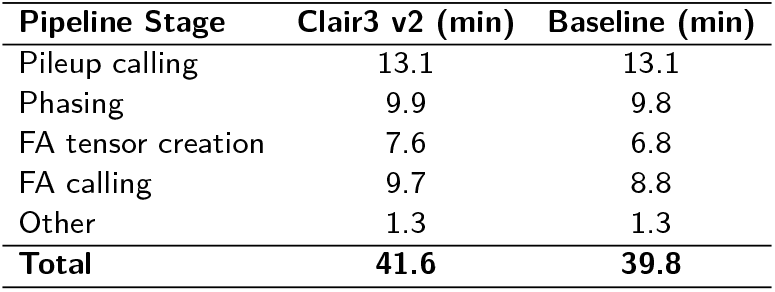
Runtime breakdown of Clair3 v2 and baseline Clair3 HAC pipeline. MV: move table features. All times are reported in minutes.

| Pipeline Stage | Clair3 v2 (min) | Baseline (min) |
| --- | --- | --- |
| Pileup calling | 13.1 | 13.1 |
| Phasing | 9.9 | 9.8 |
| FA tensor creation | 7.6 | 6.8 |
| FA calling | 9.7 | 8.8 |
| Other | 1.3 | 1.3 |
| **Total** | **41.6** | **39.8** |

Stratification by variant length revealed that Clair3 v2 preferentially improved indel detection, with minimal effect on SNPs (0.1–1.4% F1 improvement; Figure 2b,d; Supplementary Tables 5–6). For HAC basecalled reads, the benefit was most pronounced for longer indels at low coverage: at 10× depth, F1 scores for 6–15 bp insertions improved by 49.5%, and for insertions 16 bp by 37.8%. Deletions showed comparable gains, with improvements of 24.2% (6–15 bp) and 26.3% (16 bp). Although the magnitude of improvement diminished with increasing depth, substantial gains remained at 50× (e.g., 13.8% for 6–15 bp insertions). For SUP basecalled reads, the effect sizes were smaller, consistent with the reduced marginal benefit when basecalling quality is already high. The largest improvements were observed for deletions at 10× (7.6% for 6– 15 bp; 10.8% for 16 bp), whereas short insertions (1–5 bp) showed negligible or slightly negative effects at low depth.

### 3.2. Performance in complex genomic regions

To evaluate performance in challenging genomic contexts, we assessed Clair3 v2 across 16 stratified genomic regions, including low-complexity regions, homopolymers, tandem repeats, and segmental duplications, using the HG005 dataset at 30× coverage with HAC basecalling. Clair3 v2 achieved superior performance across all tested regions (Figure 3; Supplementary Table 7). We also evaluated the HG001 sample at 30× coverage and obtained similar results (Supplementary Figure 1; Supplementary Table 8).

**Figure 3.**
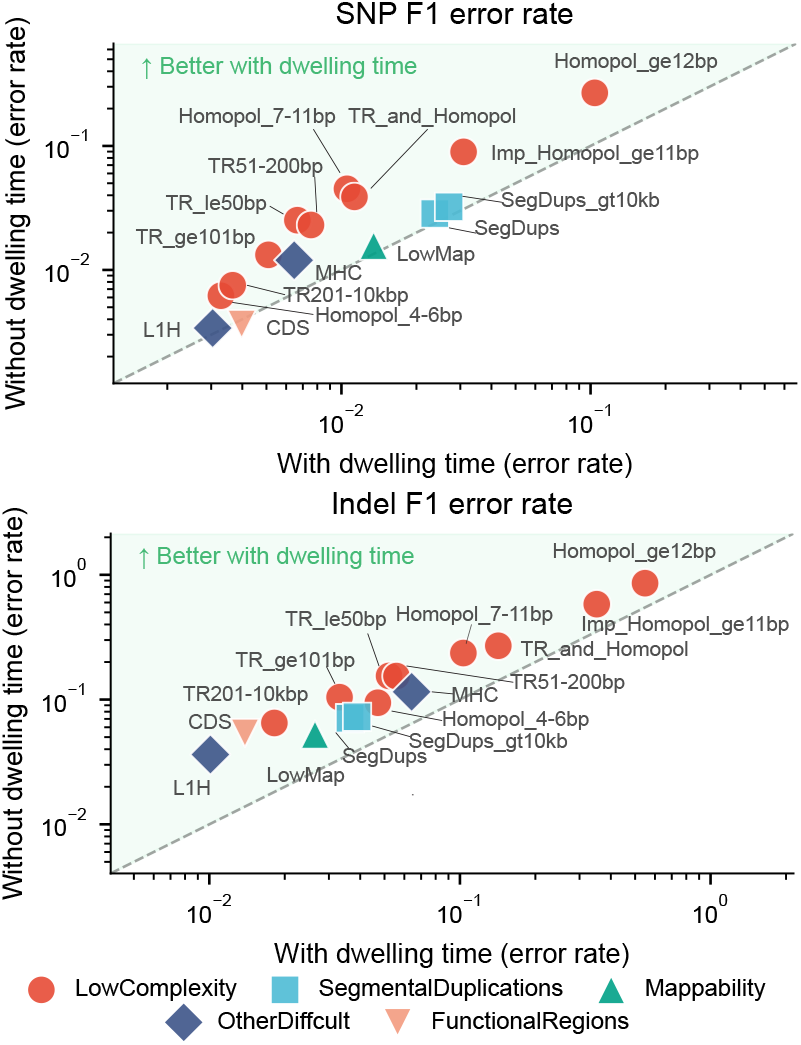
Performance in complex genomic regions. F1 error rates (1 – F1) for SNPs (top) and indels (bottom) across 16 stratified regions on HG005 at 30*×* coverage. Points above diagonal indicate improvement with Clair3 v2.

For SNP detection, dwelling time incorporation led to modest improvements in F1 score, ranging from 0.2% to 0.8% across most regions. Homopolymer-associated regions demonstrated notably larger gains: long homopolymer regions (12 bp) showed a substantial improvement from 73.2% to 89.6% F1 score, while medium-length homopolymers (7–11 bp) and imperfect homopolymers (11 bp) also benefited considerably, with F1 improvements of 3.5% and 5.9%, respectively.

For indel detection, the impact of dwelling time was more pronounced. In long homopolymer regions (12 bp), F1 score improved dramatically from 14.3% to 45.2%, while imperfect homopolymers (11 bp) rose from 41.8% to 64.9% and medium-length homopolymers (7–11 bp) from 76.4% to 89.7%. Tandem repeat regions exhibited consistent improvements ranging from 9.9% to 12.8% across different repeat length categories. For segmental duplications and low-mappability sequences, we also observed improvements of 2.6–3.4%. Importantly, Clair3 v2 also showed substantial improvements in regions excluding homopolymers ≥10 bp, with indel F1 scores improving from 93.08% to 97.55% (Supplementary Figure 2), confirming that the gains are not limited to homopolymer-rich regions.

### 3.3. Comparison with state-of-the-art variant callers

We benchmarked Clair3 v2 against DeepVariant, Dorado Variant, and LongcallD on chromosome 20 of HG003 using HAC v5.2.0 basecalling (Supplementary Table 9). This chromosome was excluded from Clair3 v2 training, and HG003 was not used in DeepVariant training, ensuring unbiased evaluation. To note, LongcallD is the only non-deep-learning method among those tested, while the currently released Dorado Variant model targets v5.0.0 basecalls, which may affect its comparative performance on v5.2.0 data.

For SNP calling, all methods achieved F1 scores greater than 99% at depths of 20× and above (Figure 4). At 10× depth, Clair3 v2 outperformed the others, achieving the highest F1 score of 98.4%, followed by DeepVariant at 97.4%.

**Figure 4.**
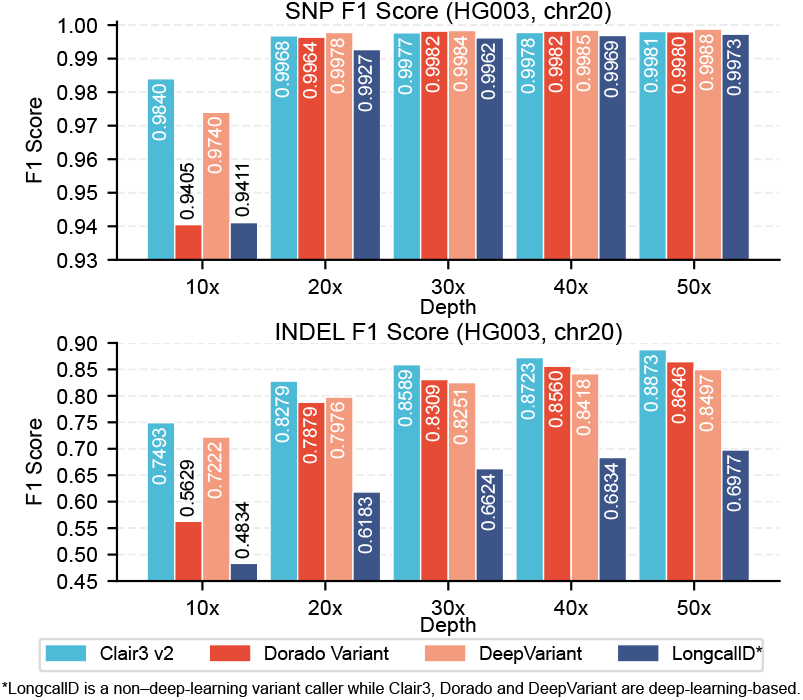
Comparison with other variant callers. SNP (top) and Indel (bottom) F1 scores on HG003 chromosome 20 at 10*×* –50*×* depth for Clair3 v2, Dorado Variant, DeepVariant, and LongcallD.

For indel calling, which proved more challenging for all methods, Clair3 v2 consistently achieved the highest F1 scores across all sequencing depths, ranging from 74.9% at 10× to 88.7% at 50×. Dorado Variant showed depth-dependent performance, lagging behind DeepVariant at lower depths (10× and 20×) but surpassing it at higher depths (30×, 40×, and 50×).

### 3.4. Computational overhead of dwelling time features

To assess the computational overhead of dwelling time features, we compared the runtime and memory usage of Clair3 v2 and baseline Clair3 using 16 threads on a machine equipped with one NVIDIA GeForce RTX 5090 D GPU and an Intel Core Ultra 7 265K processor (Tables 1 and 2). The total runtime increased marginally from 39.8 to 41.6 minutes—a difference of 1.8 minutes (4.5%). The overhead was primarily attributed to the full-alignment stages: FA tensor creation increased from 6.8 to 7.6 minutes (0.8 min overhead), and FA calling increased from 8.8 to 9.7 minutes (0.9 min overhead), demonstrating the efficiency of our genome position based circular buffer. Memory consumption remained virtually unchanged, with peak usage at 24.89 GB versus 23.60 GB and mean usage at 10.84 GB versus 10.73 GB.

**Table 2.** Memory usage of Clair3 v2 and baseline Clair3 HAC pipeline.

| Metric | Clair3 v2 (GB) | Baseline (GB) |
| --- | --- | --- |
| Peak memory usage | 24.89 | 23.60 |
| Mean memory usage | 10.84 | 10.73 |

We also benchmarked other variant callers under the same conditions using 16 threads. LongcallD, as a non-deep-learning method relying on CPU and SIMD optimizations, was the most efficient, completing in approximately 13 minutes with mean memory usage of 7.68 GB and peak memory of 27.08 GB. DeepVariant required 102 minutes with 28.5 GB mean and 52.30 GB peak memory usage. Dorado Variant, due to its whole-genome polishing approach, required substantially more computational resources: approximately 6 days on the same machine, or 2-3 hours when using a machine with 4 H800 GPUs. These results confirm that Clair3 v2 provides substantial accuracy improvements while introducing negligible computational cost, making it practical for routine use in variant calling workflows.

## 4. Discussion

We present Clair3 v2, a dwelling-time-augmented variant calling workflow that achieves substantial accuracy improvements with negligible computational overhead. Our results demonstrate that lightweight signal-level features can significantly enhance variant detection, particularly for indels in challenging genomic contexts. The most striking improvements occurred in homopolymer and tandem repeat regions, where indel F1 scores improved by up to 30%. Dwelling time provides distinct signatures for different variant types: elevated values for insertions, zero values for deletions, and unchanged values for SNPs. These patterns help the model distinguish true variants from sequencing artifacts, addressing the inherent difficulty of determining repeat lengths from basecalled sequences alone.

The magnitude of improvement varied systematically with sequencing depth and basecalling quality. At low coverage, signal features provide proportionally more value because each read carries more weight in variant calling. At higher depths, redundant coverage partially compensates for the absence of signal information. Similarly, SUP basecalling already captures substantial signal information through its sophisticated neural architecture, leaving less room for additional improvement from dwelling time features. These patterns suggest that signal-level features are most valuable in resource-constrained settings.

Compared with Dorado Variant, which also leverages move table information, Clair3 v2 achieves better performance with significantly lower computational cost. The key difference lies in approach: Dorado Variant employs whole-genome polishing that requires considerable amount of computational resources, whereas Clair3 v2 integrates signal features directly into a candidate-site-focused workflow. Our genome position based circular buffer further enhances efficiency by processing each read exactly once, eliminating redundant parsing for nearby candidate sites. This work demonstrates that signal-derived features can be effectively integrated into established variant calling pipelines, providing a practical path toward more accurate genomic inference from long-read sequencing data.

Several limitations warrant consideration. Our evaluation focused exclusively on small variants (SNPs and indels) in human genomes aligned to GRCh38; generalizability to other species, reference assemblies, and structural variants remains to be established. The quality of move-table information depends on the basecaller version, and Clair3 v2 requires BAM files generated with the move table flag, which may not be available for legacy datasets.

## 5. Conclusion

In this work, we present Clair3 v2, a signal-aware variant calling method that leverages the ONT move table to compute per-base dwelling times and integrates them as additional features in the Clair3 workflow. Through an efficient genome position based circular buffer, Clair3 v2 incorporates signal-level information with negligible computational overhead—adding less than 2 minutes to the standard pipeline runtime. Benchmarking across six GIAB samples demonstrated substantial improvements in variant calling accuracy, particularly for indels: at 10× coverage with HAC basecalling, indel F1 scores improved from 64.27% to 76.70%. The benefits were most pronounced for longer indels and in challenging genomic regions, with indel F1 scores in long homopolymer regions improving from 14.3% to 45.2%. Clair3 v2 consistently outperformed state-of-the-art variant callers including DeepVariant and Dorado Variant across all tested conditions. By demonstrating that lightweight signal-derived features can be efficiently integrated into established variant calling pipelines, this work provides a practical path toward more accurate genomic inference from long-read sequencing data.

## 8. Software and Data

Clair3 v2 is open source and available at https://github.com/HKU-BAL/Clair3/tree/clair3_v2_from_main under the BSD 3-Clause license.

All datasets used in this study are publicly available. The GRCh38 reference genome was obtained from NCBI (https://ftp.ncbi.nlm.nih.gov/genomes/all/GCA/000/001/405/GCA_000001405.15_GRCh38/). GIAB truth variant call sets (v4.2.1) for samples HG001–HG007 were downloaded from the GIAB FTP repository (https://ftp-trace.ncbi.nlm.nih.gov/giab/ftp/release/). ONT R10.4.1 (5 kHz) sequencing data for HG001–HG007 were obtained from ONT Open Data via EPI2ME (https://epi2me.nanoporetech.com/giab-2025.01/). The benchmarked commands and parameters used in this study are available in the Supplementary Materials.

## 9. Author contributions statement

R.L. conceived the study. X.Y.,Z.Z. designed the algorithms, implemented Clair3, and wrote the paper. C.L., Z.Q., M.H evaluated the benchmarking results. All authors revised the manuscript.

## 10. Acknowledgments

We thank Chris Wright, Mike Vella, Philipp Rescheneder, Sean McKenzie, Katherine Lawrence, Filipe Tostevin, and Ivan Sovic from ONT for their valuable inputs.

