## Supplementary Materials for "Leveraging ONT move table values for signal aware variant calling"

### Additional Methods and Results for Signal-Aware Variant Calling with ONT Move Tables

### Data processing procedures

#### Basecalling and alignment

Raw POD5 files were basecalled using Dorado with R10.4.1 400 bps SUP and HAC models. The human reference genome (GRCh38 no-alt analysis set) was provided during basecalling to enable reference-guided alignment. The `--emit-moves` option was enabled to output move-table information, which records the mapping between raw signal events and basecalled nucleotides. The resulting BAM files were used for downstream signal-aware variant calling experiments.

**For SUP basecalling:**

```
dorado basecaller dna_r10.4.1_e8.2_400bps_sup@v5.2.0 pod5_dir/ \
  --reference ref.fa \
  --emit-moves \
  --models-directory model_dir/ \
  > output.bam
```

**For HAC basecalling:**

```
dorado basecaller dna_r10.4.1_e8.2_400bps_hac@v5.2.0 pod5_dir/ \
  --reference ref.fa \
  --emit-moves \
  --models-directory model_dir/ \
  > output.bam
```

#### Variant calling commands

**Clair3**

```
python3 run_clair3.py \
  --bam_fn path_to_bam_file \
  --ref_fn GRCh38_no_alt_analysis_set.fasta \
  --platform ont \
  --model_path path_to_model_directory \
  --threads 60 \
  --output clair3_output \
  --sample_name SAMPLE \
  --enable_dwell_time
```

**DeepVariant**

```
singularity run deepvariant_1.9.0.sif \
  /opt/deepvariant/bin/run_deepvariant \
  --model_type ONT_R104 \
  --ref GRCh38_no_alt_analysis_set.fasta \
  --reads path_to_bam_file \
  --output_vcf deepvariant_output.vcf.gz \
  --output_gvcf deepvariant_output.g.vcf.gz \
  --num_shards 60
```

**Dorado Variant**

```
dorado variant \
  --threads 16 \
  --infer-threads 2 \
  path_to_bam_file \
  GRCh38_no_alt_analysis_set.fasta \
  --model path_to_variant_model \
  > dorado_variant_output.vcf
```

**LongcallD**

```
longcallD call \
  -t 60 \
  GRCh38_no_alt_analysis_set.fasta \
  path_to_bam_file \
  --ont \
  -n SAMPLE \
  > longcallD_output.vcf
```

#### Benchmarking commands

Variant calling performance was assessed using hap.py, qfy.py, and RTG Tools vcfeval. Predicted VCFs were compared against Genome in a Bottle high-confidence truth callsets using haplotype-aware matching with the GRCh38 reference, and, unless otherwise stated, restricted to GIAB confident regions. hap.py was run with the vcfeval engine and QUAL-based ROC generation, while qfy.py was applied to hap.py outputs to produce stratified performance reports across predefined genomic categories. In parallel, RTG Tools vcfeval was used to independently benchmark query VCFs against truth VCFs using reference SDFs. Precision, recall, and F1-score were reported for SNPs and small indels, and in selected analyses further stratified by zygosity.

**hap.py**

```
hap.py \
  path_to_truth.vcf.gz \
  path_to_query.vcf.gz \
  -o happy \
  -r GRCh38_no_alt_analysis_set.fasta \
  -f path_to_confident_regions.bed \
  --threads 70 \
  --roc QUAL \
  --engine=vcfeval \
  --engine-vcfeval-template path_to_reference_sdf
```

**qfy.py**

```
qfy.py \
  happy.vcf.gz \
  -t ga4gh \
  --stratification GRCh38-all-stratifications.tsv \
```

```
  -o strat/report \
  -r GRCh38_no_alt_analysis_set.fasta \
  --threads 60
```

**rtg tools vcfeval**

```
rtg vcfeval \
  -b path_to_truth.vcf.gz \
  -c path_to_query.vcf.gz \
  -t path_to_reference_sdf \
  --region chr20 \
  --evaluation-regions path_to_confident_regions.bed \
  --output-mode split \
  --roc-subset snp,non-snp,indel,hom,het \
  -f QUAL \
  --threads 32 \
  -o rtg_eval_output_dir
```

### Model details

#### Architecture

For the pileup model, the pileup tensor is processed sequentially using a bidirectional recurrent architecture. Two stacked bidirectional LSTM layers with hidden sizes of 128 and 160, respectively, are used to model long-range dependencies across the genomic window.The output of the recurrent layers is flattened and passed through a dropout layer (dropout rate 0.2) before branching into two task-specific fully connected heads. Each head consists of a dense layer with 128 units, predicting (i) the genotype class and (ii) the zygosity state.

For the full-alignment model, feature extraction is performed using a deep convolutional neural network composed of multiple convolutional blocks. The network applies 3 × 3 convolutions with increasing channel depth (64, 128, and 256), interleaved with ResBlocks to stabilize training and improve gradient flow. Strided convolutions (stride = 2) are used to progressively reduce spatial resolution while expanding the feature dimension. To aggregate spatial information across varying read depths and local contexts, the network incorporates a spatial pyramid pooling (SPP) layer, which pools feature maps at multiple spatial scales (e.g., 1 × 1, 2 × 2, and 3 × 3).After pyramid pooling, features are flattened and regularized using dropout (dropout rate 0.2), followed by a dense layer with 256 units. The model then branches into four parallel prediction heads, each implemented as a dense layer with 128 units, to predict genotype, zygosity, and two indel length–related outputs. Additional dropout (0.5 and 0.2) is applied in the prediction heads to mitigate overfitting.

#### Description of DNA pileup input features

The pileup tensor has shape [33, 18], representing a 33-position window (flankingBaseNum=16) centered on the candidate variant. The 18 feature channels encode strand-specific nucleotide counts and indel information: A, C, G, T, I, I1, D, D1, *, a, c, g, t, i, i1, d, d1, #. Channels follow a strand-specific encoding scheme where uppercase letters represent one strand and lowercase letters represent the complementary strand:

1. Base count channels (A/a, C/c, G/g, T/t): Strand-specific counts of each nucleotide at each position. These channels capture the primary sequence information and strand bias patterns.
2. Insertion channels (I/i, I1/i1): I/i channels represent the total insertion support at each position (summed across all insertion variants), while I1/i1 channels specifically count the most frequently observed insertion variant. This dual representation allows the model to distinguish between positions with a single dominant insertion versus positions with multiple competing insertions.
3. Deletion channels (D/d, D1/d1): D/d channels represent the total support for deletions starting at each position, while D1/d1 channels count only the most frequently observed deletion. Similar to insertions, this enables detection of deletion variants and assessment of their complexity.
4. Deletion continuation channels (*, #): These channels mark positions that are within a deletion span but not the deletion start position. This allows the model to distinguish between deletion-start positions and deletion-continuation positions.
5. Reference encoding: Reference base channels are encoded using a signed representation where the reference base position is overwritten with the negative sum of the opposite strand's base counts. This encoding explicitly marks the reference base while preserving strand-specific information.

#### Description of DNA full-alignment input features

The full-alignment tensor has shape [89, 33, C] where 89 represents the maximum read depth for ONT data, 33 represents genomic positions in the window around the candidate variant, and C is the number of feature channels (8 in standard mode, 9 when dwell-time is enabled). Reads are sorted by haplotype tag and read order before tensor construction. When coverage exceeds 89 reads, excess reads are downsampled to maintain the fixed tensor dimensions.

1. Reference base: Encoded representation of the reference genome base at each position (A, C, G, T mapped to fixed numeric codes).
2. Alternative base: Encodes the read base at each position, specifically highlighting mismatches and indel types relative to the reference. This channel captures variant evidence at the read level.
3. Strand information: Binary encoding indicating whether each read is mapped to the forward or reverse strand, enabling detection of strand-specific artifacts and true strand bias associated with variants.
4. Mapping quality: Per-read mapping quality normalized to the range [0, 100], with values capped at MQ=60. This channel helps the model downweight poorly aligned reads.
5. Base quality: Per-base quality score normalized to [0, 100], with values capped at BQ=40. This provides position-specific confidence in the sequenced nucleotide.
6. Variant support: Normalized per-read allele frequency support for the candidate variant, scaled to [0, 100]. This channel explicitly marks which reads support the variant versus the reference allele.

7. Inserted bases: Encodes insertion sequences by projecting the inserted bases across subsequent positions in the read. This representation allows the convolutional layers to process insertion content spatially.
8. Phasing information: Encodes haplotype assignments (haplotype 1, haplotype 2, or unphased) derived from BAM HP tags. This enables the model to leverage phasing information when available, improving accuracy for compound heterozygous variants and linked polymorphisms.
9. Dwelling time (enabled by `--enable_dwell_time`): This channel is extracted from the ONT BAM mv tag (move table) and aligned to read positions. Dwell-time features provide nanopore-specific signal-level information that can improve variant discrimination, particularly for homopolymers and low-complexity regions where base-calling uncertainty is high.

#### Description of network outputs

Clair3 is formulated as a multi-task classification problem with up to four prediction heads, enabling simultaneous prediction of variant genotype, variant type, and structural characteristics:

- **GT21 head (21 classes):** Predicts the specific genotype combination from 21 possible classes: AA, AC, AG, AT, CC, CG, CT, GG, GT, TT (SNV genotypes), DelDel, ADel, CDel, GDel, TDel (deletion genotypes), InsIns, AIns, CIns, GIns, TIns (insertion genotypes), and InsDel (compound indel). This fine-grained classification enables precise variant representation.
- **Genotype class head (3 classes):** Classifies variants into three zygosity states: 0/0 (homozygous reference), 1/1 (homozygous variant), and 0/1 (heterozygous variant, which includes both 0/1 and 1/2 genotypes merged into a single heterozygous class).
- **Indel length heads (2× 33 classes):** For the full-alignment model, two additional heads predict the length of insertions or deletions for each allele. Each head outputs a 33 class softmax distribution representing indel lengths from -16 (16 bp deletion) to +16 (16 bp insertion), with lengths clipped to this range.

#### Training configurations

Models were trained using distributed data-parallel execution across four GPUs. Optimization was performed with the AdamW optimizer and a base learning rate of $1 \times 10^{-4}$. Learning-rate scaling was enabled, such that the effective learning rate was multiplied by the number of GPUs; with four GPUs, this resulted in an effective learning rate of $4 \times 10^{-4}$.

Training was conducted for a maximum of 30 epochs. Early stopping was applied based on validation loss, with a patience of 10 epochs. The checkpoint corresponding to the lowest validation loss was retained for downstream evaluation.

A random split of the training data was used to form training and validation sets, with 90% of samples used for training and 10% for validation. The loss function consisted of per-head focal loss (focusing parameter $\gamma = 2$), summed across all active prediction heads. L2 weight decay for full-alignment training was set to $1 \times 10^{-7}$.

### Supplementary tables

***Supplementary table 1*** *Clair3 v2 (HAC) performance*

| Type | Sample | TRUTH.TOTAL | Depth | TRUTH.TP | TRUTH.FN | QUERY.TOTAL | QUERY.FP | QUERY.UNK | FP.gt | FP.al | METRIC.Recall | METRIC.Precision | METRIC.F1_Score |
|---|---|---|---|---|---|---|---|---|---|---|---|---|---|
| SNP | HG001 | 3,254,386 | 10x | 3,138,888 | 115,498 | 4,085,979 | 41,887 | 904,602 | 26,189 | 3,572 | 96.45% | 98.68% | 97.55% |
| | | | 20x | 3,232,222 | 22,164 | 4,279,492 | 9,420 | 1,036,931 | 1,982 | 1,684 | 99.32% | 99.71% | 99.51% |
| | | | 30x | 3,240,024 | 14,362 | 4,378,287 | 6,074 | 1,131,143 | 863 | 1,102 | 99.56% | 99.81% | 99.69% |
| | | | 40x | 3,244,675 | 9,711 | 4,435,819 | 5,362 | 1,184,632 | 724 | 862 | 99.70% | 99.84% | 99.77% |
| | | | 50x | 3,247,795 | 6,591 | 4,484,603 | 4,562 | 1,231,073 | 710 | 720 | 99.80% | 99.86% | 99.83% |
| | HG003 | 3,327,495 | 10x | 3,218,821 | 108,674 | 4,144,864 | 39,226 | 885,923 | 24,157 | 3,879 | 96.73% | 98.80% | 97.75% |
| | | | 20x | 3,297,399 | 30,096 | 4,343,070 | 10,541 | 1,033,832 | 2,540 | 2,214 | 99.10% | 99.68% | 99.39% |
| | | | 30x | 3,304,347 | 23,148 | 4,443,503 | 7,171 | 1,130,507 | 1,384 | 1,398 | 99.30% | 99.78% | 99.54% |
| | | | 40x | 3,309,611 | 17,884 | 4,492,382 | 6,222 | 1,175,022 | 1,203 | 1,161 | 99.46% | 99.81% | 99.64% |
| | | | 50x | 3,313,946 | 13,549 | 4,537,053 | 5,888 | 1,215,635 | 1,281 | 1,061 | 99.59% | 99.82% | 99.71% |
| | HG004 | 3,346,610 | 10x | 3,225,924 | 120,686 | 4,144,406 | 40,469 | 877,052 | 27,463 | 3,805 | 96.39% | 98.76% | 97.56% |
| | | | 20x | 3,315,702 | 30,908 | 4,360,400 | 8,849 | 1,034,419 | 2,525 | 1,863 | 99.08% | 99.73% | 99.40% |
| | | | 30x | 3,324,269 | 22,341 | 4,460,165 | 6,141 | 1,128,217 | 1,300 | 1,444 | 99.33% | 99.82% | 99.57% |
| | | | 40x | 3,330,232 | 16,378 | 4,514,894 | 5,181 | 1,177,901 | 1,272 | 1,208 | 99.51% | 99.84% | 99.68% |
| | | | 50x | 3,334,931 | 11,679 | 4,563,123 | 4,120 | 1,222,428 | 1,167 | 1,054 | 99.65% | 99.88% | 99.76% |
| | HG005 | 3,275,631 | 10x | 3,170,659 | 104,972 | 4,113,268 | 39,149 | 902,690 | 23,492 | 3,516 | 96.80% | 98.78% | 97.78% |
| | | | 20x | 3,252,004 | 23,627 | 4,306,863 | 8,555 | 1,045,183 | 2,368 | 1,497 | 99.28% | 99.74% | 99.51% |
| | | | 30x | 3,260,071 | 15,560 | 4,420,498 | 6,120 | 1,153,037 | 1,003 | 1,052 | 99.53% | 99.81% | 99.67% |
| | | | 40x | 3,264,268 | 11,363 | 4,475,368 | 5,499 | 1,204,291 | 839 | 910 | 99.65% | 99.83% | 99.74% |
| | | | 50x | 3,267,558 | 8,073 | 4,521,883 | 4,450 | 1,248,537 | 821 | 801 | 99.75% | 99.86% | 99.81% |
| | HG006 | 3,269,860 | 10x | 3,165,484 | 104,376 | 4,072,592 | 39,430 | 867,195 | 24,506 | 2,881 | 96.81% | 98.77% | 97.78% |
| | | | 20x | 3,244,483 | 25,377 | 4,264,902 | 9,141 | 1,010,576 | 2,563 | 1,226 | 99.22% | 99.72% | 99.47% |
| | | | 30x | 3,251,550 | 18,310 | 4,370,470 | 6,382 | 1,111,738 | 1,207 | 869 | 99.44% | 99.80% | 99.62% |
| | | | 40x | 3,256,211 | 13,649 | 4,442,085 | 6,411 | 1,178,644 | 1,169 | 797 | 99.58% | 99.80% | 99.69% |
| | | | 50x | 3,260,086 | 9,774 | 4,487,962 | 5,506 | 1,221,540 | 1,080 | 680 | 99.70% | 99.83% | 99.77% |
| | HG007 | 3,284,462 | 10x | 3,173,896 | 110,566 | 4,090,800 | 39,029 | 877,408 | 26,354 | 2,696 | 96.63% | 98.79% | 97.70% |
| | | | 20x | 3,252,281 | 32,181 | 4,279,621 | 8,530 | 1,018,119 | 2,923 | 1,246 | 99.02% | 99.74% | 99.38% |
| | | | 30x | 3,260,661 | 23,801 | 4,368,895 | 5,622 | 1,101,857 | 1,237 | 870 | 99.28% | 99.83% | 99.55% |
| | | | 40x | 3,265,598 | 18,864 | 4,429,877 | 4,788 | 1,158,700 | 1,094 | 726 | 99.43% | 99.85% | 99.64% |
| | | | 50x | 3,271,127 | 13,335 | 4,489,217 | 4,613 | 1,212,684 | 1,075 | 747 | 99.59% | 99.86% | 99.73% |
| INDEL | HG001 | 467,702 | 10x | 294,222 | 173,480 | 527,091 | 28,829 | 198,324 | 11,982 | 9,569 | 62.91% | 91.23% | 74.47% |
| | | | 20x | 351,018 | 116,684 | 632,251 | 24,325 | 248,644 | 8,917 | 10,794 | 75.05% | 93.66% | 83.33% |
| | | | 30x | 372,097 | 95,605 | 680,950 | 24,156 | 275,308 | 8,575 | 11,645 | 79.56% | 94.05% | 86.20% |
| | | | 40x | 384,154 | 83,548 | 711,806 | 24,281 | 293,251 | 8,666 | 12,137 | 82.14% | 94.20% | 87.76% |
| | | | 50x | 391,619 | 76,083 | 732,041 | 24,305 | 305,484 | 8,756 | 12,493 | 83.73% | 94.30% | 88.70% |
| | HG003 | 504,501 | 10x | 317,355 | 187,146 | 545,663 | 30,084 | 190,871 | 12,578 | 10,435 | 62.90% | 91.52% | 74.56% |
| | | | 20x | 373,284 | 131,217 | 644,975 | 25,638 | 235,809 | 9,434 | 11,631 | 73.99% | 93.73% | 82.70% |
| | | | 30x | 395,198 | 109,303 | 692,424 | 25,769 | 259,975 | 9,135 | 12,848 | 78.33% | 94.04% | 85.47% |
| | | | 40x | 407,634 | 96,867 | 722,948 | 26,519 | 276,614 | 9,500 | 13,761 | 80.80% | 94.06% | 86.93% |
| | | | 50x | 415,558 | 88,943 | 743,228 | 26,836 | 288,220 | 9,813 | 14,125 | 82.37% | 94.10% | 87.85% |
| | HG004 | 510,519 | 10x | 315,798 | 194,721 | 542,162 | 30,510 | 188,642 | 12,920 | 10,641 | 61.86% | 91.37% | 73.77% |
| | | | 20x | 374,443 | 136,076 | 645,439 | 25,775 | 235,058 | 9,317 | 11,954 | 73.35% | 93.72% | 82.29% |
| | | | 30x | 396,487 | 114,032 | 691,800 | 26,045 | 257,858 | 9,124 | 13,164 | 77.66% | 94.00% | 85.05% |
| | | | 40x | 408,827 | 101,692 | 721,084 | 26,821 | 273,287 | 9,464 | 14,232 | 80.08% | 94.01% | 86.49% |
| | | | 50x | 417,129 | 93,390 | 742,286 | 27,190 | 285,330 | 9,732 | 14,669 | 81.71% | 94.05% | 87.44% |
| | HG005 | 416,777 | 10x | 301,753 | 115,024 | 536,421 | 26,229 | 202,086 | 11,486 | 8,150 | 72.40% | 92.15% | 81.09% |
| | | | 20x | 352,016 | 64,761 | 637,290 | 18,930 | 257,455 | 7,769 | 7,294 | 84.46% | 95.02% | 89.43% |
| | | | 30x | 368,605 | 48,172 | 685,577 | 16,279 | 290,810 | 6,541 | 6,736 | 88.44% | 95.88% | 92.01% |
| | | | 40x | 376,878 | 39,899 | 716,865 | 14,750 | 314,838 | 5,911 | 6,398 | 90.43% | 96.33% | 93.29% |
| | | | 50x | 381,528 | 35,249 | 736,786 | 13,681 | 330,861 | 5,581 | 6,088 | 91.54% | 96.63% | 94.02% |
| | HG006 | 428,237 | 10x | 291,574 | 136,663 | 542,029 | 24,735 | 219,417 | 10,391 | 7,913 | 68.09% | 92.33% | 78.38% |
| | | | 20x | 338,852 | 89,386 | 641,810 | 19,255 | 274,958 | 6,844 | 8,238 | 79.13% | 94.75% | 86.24% |
| | | | 30x | 356,052 | 72,186 | 690,179 | 18,007 | 306,408 | 5,901 | 8,674 | 83.14% | 95.31% | 88.81% |
| | | | 40x | 364,895 | 63,343 | 720,816 | 17,861 | 327,886 | 5,857 | 9,021 | 85.21% | 95.45% | 90.04% |
| | | | 50x | 370,557 | 57,681 | 741,980 | 17,739 | 343,176 | 5,976 | 9,134 | 86.53% | 95.55% | 90.82% |
| | HG007 | 430,326 | 10x | 290,345 | 139,981 | 542,676 | 25,071 | 220,852 | 10,698 | 7,832 | 67.47% | 92.21% | 77.92% |
| | | | 20x | 337,410 | 92,916 | 642,393 | 19,808 | 276,459 | 7,159 | 8,475 | 78.41% | 94.59% | 85.74% |
| | | | 30x | 354,379 | 75,947 | 689,777 | 18,736 | 307,015 | 6,258 | 9,204 | 82.35% | 95.11% | 88.27% |
| | | | 40x | 362,983 | 67,343 | 719,379 | 18,517 | 327,742 | 6,248 | 9,510 | 84.35% | 95.27% | 89.48% |
| | | | 50x | 361,777 | 68,549 | 733,353 | 28,926 | 332,296 | 7,639 | 10,945 | 84.07% | 92.79% | 88.21% |

***Supplementary table 2*** *Clair3 baseline (HAC) performance*

| Type | Sample | TRUTH.TOTAL | Depth | TRUTH.TP | TRUTH.FN | QUERY.TOTAL | QUERY.FP | QUERY.UNK | FP.gt | FP.al | METRIC.Recall | METRIC.Precision | METRIC.F1_Score |
|---|---|---|---|---|---|---|---|---|---|---|---|---|---|
| SNP | HG001 | 3,254,386 | 10x | 3,051,760 | 202,626 | 3,825,515 | 46,054 | 727,428 | 31,562 | 2,705 | 93.77% | 98.51% | 96.09% |
| | | | 20x | 3,214,684 | 39,702 | 4,024,045 | 14,043 | 794,970 | 6,946 | 1,171 | 98.78% | 99.57% | 99.17% |
| | | | 30x | 3,233,756 | 20,630 | 4,076,642 | 9,576 | 832,967 | 4,915 | 610 | 99.37% | 99.70% | 99.54% |
| | | | 40x | 3,239,150 | 15,236 | 4,100,000 | 8,765 | 851,717 | 4,342 | 502 | 99.53% | 99.73% | 99.63% |
| | | | 50x | 3,241,071 | 13,315 | 4,118,035 | 7,874 | 868,754 | 4,150 | 441 | 99.59% | 99.76% | 99.67% |
| | HG003 | 3,327,495 | 10x | 3,152,420 | 175,075 | 3,892,164 | 41,750 | 697,464 | 27,825 | 3,031 | 94.74% | 98.69% | 96.68% |
| | | | 20x | 3,288,252 | 39,243 | 4,072,741 | 14,959 | 768,889 | 6,880 | 1,741 | 98.82% | 99.55% | 99.18% |
| | | | 30x | 3,302,908 | 24,587 | 4,125,233 | 11,419 | 810,257 | 5,421 | 1,032 | 99.26% | 99.66% | 99.46% |
| | | | 40x | 3,306,474 | 21,021 | 4,145,060 | 9,844 | 828,087 | 4,989 | 728 | 99.37% | 99.70% | 99.54% |
| | | | 50x | 3,307,846 | 19,649 | 4,158,599 | 9,344 | 840,767 | 4,824 | 665 | 99.41% | 99.72% | 99.56% |
| | HG004 | 3,346,610 | 10x | 3,155,219 | 191,391 | 3,892,149 | 41,714 | 694,634 | 30,149 | 2,929 | 94.28% | 98.70% | 96.44% |
| | | | 20x | 3,306,733 | 39,877 | 4,093,281 | 13,523 | 772,299 | 6,713 | 1,506 | 98.81% | 99.59% | 99.20% |
| | | | 30x | 3,323,749 | 22,861 | 4,147,109 | 9,730 | 812,945 | 5,071 | 983 | 99.32% | 99.71% | 99.51% |
| | | | 40x | 3,328,379 | 18,231 | 4,166,347 | 8,492 | 828,758 | 4,641 | 780 | 99.46% | 99.75% | 99.60% |
| | | | 50x | 3,329,999 | 16,611 | 4,188,795 | 8,023 | 850,066 | 4,433 | 781 | 99.50% | 99.76% | 99.63% |
| | HG005 | 3,275,631 | 10x | 3,091,027 | 184,604 | 3,863,569 | 46,024 | 726,089 | 31,093 | 2,748 | 94.36% | 98.53% | 96.40% |
| | | | 20x | 3,237,317 | 38,314 | 4,044,217 | 13,915 | 792,508 | 7,338 | 1,218 | 98.83% | 99.57% | 99.20% |
| | | | 30x | 3,255,054 | 20,577 | 4,105,183 | 10,938 | 838,658 | 5,296 | 891 | 99.37% | 99.67% | 99.52% |
| | | | 40x | 3,259,643 | 15,988 | 4,121,060 | 9,829 | 851,038 | 4,909 | 664 | 99.51% | 99.70% | 99.61% |
| | | | 50x | 3,261,167 | 14,464 | 4,136,162 | 8,750 | 865,704 | 4,831 | 604 | 99.56% | 99.73% | 99.65% |
| | HG006 | 3,269,860 | 10x | 3,094,067 | 175,793 | 3,837,217 | 43,916 | 698,931 | 30,689 | 2,381 | 94.62% | 98.60% | 96.57% |
| | | | 20x | 3,233,864 | 35,996 | 4,017,273 | 14,480 | 768,601 | 7,386 | 1,018 | 98.90% | 99.55% | 99.23% |
| | | | 30x | 3,249,729 | 20,131 | 4,066,130 | 10,543 | 805,549 | 5,178 | 591 | 99.38% | 99.68% | 99.53% |
| | | | 40x | 3,253,770 | 16,090 | 4,096,668 | 10,454 | 832,154 | 4,903 | 534 | 99.51% | 99.68% | 99.59% |
| | | | 50x | 3,255,118 | 14,742 | 4,109,476 | 9,479 | 844,594 | 4,676 | 450 | 99.55% | 99.71% | 99.63% |
| | HG007 | 3,284,462 | 10x | 3,105,808 | 178,654 | 3,858,409 | 44,171 | 708,175 | 32,486 | 2,265 | 94.56% | 98.60% | 96.54% |
| | | | 20x | 3,244,677 | 39,785 | 4,034,654 | 13,544 | 776,110 | 7,720 | 943 | 98.79% | 99.58% | 99.18% |
| | | | 30x | 3,261,365 | 23,097 | 4,076,178 | 9,891 | 804,661 | 5,369 | 599 | 99.30% | 99.70% | 99.50% |
| | | | 40x | 3,265,430 | 19,032 | 4,099,790 | 8,517 | 825,572 | 4,877 | 473 | 99.42% | 99.74% | 99.58% |
| | | | 50x | 3,265,436 | 19,026 | 4,115,213 | 9,169 | 840,365 | 5,231 | 505 | 99.42% | 99.72% | 99.57% |
| INDEL | HG001 | 467,702 | 10x | 208,617 | 259,085 | 313,686 | 8,201 | 95,064 | 4,334 | 1,342 | 44.60% | 96.25% | 60.96% |
| | | | 20x | 284,022 | 183,680 | 435,781 | 7,212 | 140,377 | 3,972 | 1,835 | 60.73% | 97.56% | 74.86% |
| | | | 30x | 309,962 | 157,740 | 482,710 | 7,148 | 159,893 | 4,125 | 2,170 | 66.27% | 97.79% | 79.00% |
| | | | 40x | 323,307 | 144,395 | 507,788 | 7,021 | 170,936 | 4,128 | 2,301 | 69.13% | 97.92% | 81.04% |
| | | | 50x | 329,985 | 137,717 | 520,063 | 6,696 | 176,470 | 4,007 | 2,237 | 70.55% | 98.05% | 82.06% |
| | HG003 | 504,501 | 10x | 232,475 | 272,026 | 340,145 | 8,864 | 96,265 | 4,700 | 1,534 | 46.08% | 96.37% | 62.35% |
| | | | 20x | 307,241 | 197,260 | 458,435 | 7,460 | 138,109 | 4,126 | 2,048 | 60.90% | 97.67% | 75.02% |
| | | | 30x | 332,203 | 172,298 | 502,058 | 7,279 | 155,307 | 4,221 | 2,304 | 65.85% | 97.90% | 78.74% |
| | | | 40x | 344,684 | 159,817 | 524,714 | 7,312 | 164,512 | 4,317 | 2,494 | 68.32% | 97.97% | 80.50% |
| | | | 50x | 351,041 | 153,460 | 535,636 | 7,219 | 168,803 | 4,350 | 2,493 | 69.58% | 98.03% | 81.39% |
| | HG004 | 510,519 | 10x | 229,250 | 281,269 | 334,233 | 9,204 | 93,441 | 4,872 | 1,570 | 44.91% | 96.18% | 61.22% |
| | | | 20x | 306,699 | 203,820 | 456,826 | 7,521 | 137,199 | 4,087 | 2,043 | 60.08% | 97.65% | 74.39% |
| | | | 30x | 332,299 | 178,220 | 501,053 | 7,189 | 154,416 | 4,044 | 2,354 | 65.09% | 97.93% | 78.20% |
| | | | 40x | 345,704 | 164,815 | 525,462 | 7,206 | 164,437 | 4,134 | 2,619 | 67.72% | 98.00% | 80.09% |
| | | | 50x | 352,657 | 157,862 | 537,838 | 7,090 | 169,535 | 4,101 | 2,612 | 69.08% | 98.08% | 81.06% |
| | HG005 | 416,777 | 10x | 219,075 | 197,702 | 325,872 | 8,151 | 96,531 | 4,469 | 1,187 | 52.56% | 96.45% | 68.04% |
| | | | 20x | 290,561 | 126,216 | 441,969 | 7,008 | 139,822 | 4,101 | 1,575 | 69.72% | 97.68% | 81.36% |
| | | | 30x | 315,055 | 101,722 | 487,220 | 6,531 | 159,438 | 4,078 | 1,649 | 75.59% | 98.01% | 85.35% |
| | | | 40x | 326,927 | 89,850 | 510,423 | 6,504 | 170,061 | 4,225 | 1,700 | 78.44% | 98.09% | 87.17% |
| | | | 50x | 332,963 | 83,814 | 521,865 | 6,306 | 175,299 | 4,169 | 1,675 | 79.89% | 98.18% | 88.10% |
| | HG006 | 428,237 | 10x | 217,979 | 210,258 | 337,112 | 7,976 | 109,004 | 4,246 | 1,269 | 50.90% | 96.50% | 66.65% |
| | | | 20x | 285,696 | 142,542 | 452,691 | 6,552 | 155,739 | 3,635 | 1,615 | 66.71% | 97.79% | 79.32% |
| | | | 30x | 307,749 | 120,489 | 496,132 | 6,045 | 176,066 | 3,454 | 1,827 | 71.86% | 98.11% | 82.96% |
| | | | 40x | 318,369 | 109,869 | 519,132 | 6,001 | 187,664 | 3,580 | 1,840 | 74.34% | 98.19% | 84.62% |
| | | | 50x | 323,650 | 104,588 | 530,099 | 5,844 | 193,162 | 3,518 | 1,840 | 75.58% | 98.27% | 85.44% |
| | HG007 | 430,326 | 10x | 217,855 | 212,471 | 338,964 | 8,187 | 110,688 | 4,245 | 1,286 | 50.63% | 96.41% | 66.39% |
| | | | 20x | 285,101 | 145,225 | 454,794 | 6,703 | 158,164 | 3,668 | 1,725 | 66.25% | 97.74% | 78.97% |
| | | | 30x | 306,633 | 123,693 | 496,599 | 6,029 | 177,631 | 3,514 | 1,862 | 71.26% | 98.11% | 82.55% |
| | | | 40x | 317,082 | 113,244 | 519,539 | 6,137 | 189,264 | 3,634 | 2,050 | 73.68% | 98.14% | 84.17% |
| | | | 50x | 317,105 | 113,221 | 525,761 | 8,548 | 192,765 | 4,540 | 2,558 | 73.69% | 97.43% | 83.91% |

#### ***Supplementary table 3*** *Clair3 v2 (SUP) performance*

| Type | Sample | TRUTH.TOTAL | Depth | TRUTH.TP | TRUTH.FN | QUERY.TOTAL | QUERY.FP | QUERY.UNK | FP.gt | FP.al | METRIC.Recall | METRIC.Precision | METRIC.F1_Score |
|---|---|---|---|---|---|---|---|---|---|---|---|---|---|
| SNP | HG001 | 3,254,386 | 10x | 3,167,968 | 86,418 | 4,051,017 | 33,142 | 849,191 | 20,166 | 2,055 | 97.34% | 98.96% | 98.15% |
| | | | 20x | 3,241,958 | 12,428 | 4,210,748 | 7,566 | 960,129 | 1,385 | 907 | 99.62% | 99.77% | 99.69% |
| | | | 30x | 3,247,237 | 7,149 | 4,276,544 | 5,521 | 1,022,574 | 787 | 605 | 99.78% | 99.83% | 99.81% |
| | | | 40x | 3,249,447 | 4,939 | 4,326,246 | 5,348 | 1,070,208 | 777 | 547 | 99.85% | 99.84% | 99.84% |
| | | | 50x | 3,250,704 | 3,682 | 4,365,929 | 4,245 | 1,109,676 | 724 | 441 | 99.89% | 99.87% | 99.88% |
| | HG003 | 3,327,495 | 10x | 3,246,353 | 81,142 | 4,104,739 | 32,993 | 824,408 | 19,429 | 2,817 | 97.56% | 98.99% | 98.27% |
| | | | 20x | 3,308,951 | 18,544 | 4,259,071 | 8,327 | 940,347 | 1,708 | 1,333 | 99.44% | 99.75% | 99.60% |
| | | | 30x | 3,313,597 | 13,898 | 4,332,755 | 6,003 | 1,011,545 | 1,146 | 851 | 99.58% | 99.82% | 99.70% |
| | | | 40x | 3,315,994 | 11,501 | 4,383,838 | 5,324 | 1,060,846 | 1,132 | 757 | 99.65% | 99.84% | 99.75% |
| | | | 50x | 3,317,620 | 9,875 | 4,428,419 | 4,947 | 1,104,142 | 1,155 | 712 | 99.70% | 99.85% | 99.78% |
| | HG004 | 3,346,610 | 10x | 3,257,583 | 89,027 | 4,110,636 | 31,737 | 820,249 | 20,981 | 2,387 | 97.34% | 99.04% | 98.18% |
| | | | 20x | 3,328,447 | 18,163 | 4,270,054 | 7,320 | 932,776 | 1,723 | 1,175 | 99.46% | 99.78% | 99.62% |
| | | | 30x | 3,334,059 | 12,551 | 4,346,468 | 4,987 | 1,005,763 | 1,108 | 840 | 99.63% | 99.85% | 99.74% |
| | | | 40x | 3,337,198 | 9,412 | 4,400,495 | 4,166 | 1,057,418 | 1,101 | 744 | 99.72% | 99.88% | 99.80% |
| | | | 50x | 3,338,935 | 7,675 | 4,443,723 | 3,588 | 1,099,460 | 1,105 | 720 | 99.77% | 99.89% | 99.83% |
| | HG005 | 3,275,631 | 10x | 3,198,969 | 76,662 | 4,082,361 | 31,612 | 850,900 | 18,414 | 2,160 | 97.66% | 99.02% | 98.34% |
| | | | 20x | 3,261,567 | 14,064 | 4,227,522 | 7,201 | 957,500 | 1,587 | 953 | 99.57% | 99.78% | 99.68% |
| | | | 30x | 3,267,179 | 8,452 | 4,306,302 | 5,150 | 1,032,637 | 825 | 704 | 99.74% | 99.84% | 99.79% |
| | | | 40x | 3,269,297 | 6,334 | 4,363,187 | 4,667 | 1,087,806 | 804 | 618 | 99.81% | 99.86% | 99.83% |
| | | | 50x | 3,270,410 | 5,221 | 4,402,352 | 4,001 | 1,126,493 | 821 | 574 | 99.84% | 99.88% | 99.86% |
| | HG006 | 3,269,860 | 10x | 3,192,027 | 77,833 | 4,042,617 | 34,285 | 815,771 | 19,564 | 1,934 | 97.62% | 98.94% | 98.27% |
| | | | 20x | 3,254,894 | 14,966 | 4,184,207 | 7,771 | 920,762 | 1,756 | 781 | 99.54% | 99.76% | 99.65% |
| | | | 30x | 3,259,919 | 9,941 | 4,266,016 | 5,388 | 999,881 | 999 | 599 | 99.70% | 99.84% | 99.77% |
| | | | 40x | 3,262,368 | 7,492 | 4,323,625 | 4,976 | 1,055,429 | 995 | 567 | 99.77% | 99.85% | 99.81% |
| | | | 50x | 3,263,603 | 6,257 | 4,368,613 | 4,629 | 1,099,507 | 1,005 | 502 | 99.81% | 99.86% | 99.83% |
| | HG007 | 3,284,462 | 10x | 3,203,440 | 81,022 | 4,056,765 | 31,987 | 820,863 | 20,780 | 1,812 | 97.53% | 99.01% | 98.27% |
| | | | 20x | 3,266,102 | 18,360 | 4,192,091 | 6,919 | 918,369 | 1,812 | 834 | 99.44% | 99.79% | 99.61% |
| | | | 30x | 3,271,723 | 12,739 | 4,261,007 | 4,796 | 983,702 | 984 | 591 | 99.61% | 99.85% | 99.73% |
| | | | 40x | 3,274,449 | 10,013 | 4,317,706 | 4,411 | 1,038,022 | 953 | 501 | 99.70% | 99.87% | 99.78% |
| | | | 50x | 3,276,269 | 8,193 | 4,363,558 | 3,957 | 1,082,500 | 970 | 474 | 99.75% | 99.88% | 99.81% |
| INDEL | HG001 | 467,702 | 10x | 312,437 | 155,265 | 559,207 | 25,088 | 215,536 | 12,377 | 7,792 | 66.80% | 92.70% | 77.65% |
| | | | 20x | 369,307 | 98,395 | 673,181 | 23,668 | 271,040 | 10,240 | 9,552 | 78.96% | 94.11% | 85.88% |
| | | | 30x | 388,730 | 78,972 | 719,059 | 24,023 | 295,876 | 10,103 | 10,682 | 83.11% | 94.32% | 88.37% |
| | | | 40x | 399,549 | 68,153 | 746,592 | 23,703 | 312,104 | 9,845 | 11,020 | 85.43% | 94.54% | 89.76% |
| | | | 50x | 406,365 | 61,337 | 764,322 | 23,550 | 322,748 | 9,760 | 11,276 | 86.89% | 94.67% | 90.61% |
| | HG003 | 504,501 | 10x | 336,949 | 167,552 | 581,511 | 27,643 | 208,952 | 13,688 | 8,676 | 66.79% | 92.58% | 77.60% |
| | | | 20x | 393,182 | 111,319 | 686,209 | 25,809 | 256,139 | 11,359 | 10,643 | 77.93% | 94.00% | 85.22% |
| | | | 30x | 413,546 | 90,955 | 729,390 | 25,848 | 277,585 | 10,992 | 11,782 | 81.97% | 94.28% | 87.70% |
| | | | 40x | 424,511 | 79,990 | 754,718 | 25,639 | 291,354 | 10,786 | 12,209 | 84.14% | 94.47% | 89.01% |
| | | | 50x | 431,789 | 72,712 | 771,567 | 25,677 | 300,515 | 10,798 | 12,420 | 85.59% | 94.55% | 89.85% |
| | HG004 | 510,519 | 10x | 335,872 | 174,647 | 577,438 | 27,842 | 205,900 | 13,929 | 8,802 | 65.79% | 92.51% | 76.89% |
| | | | 20x | 393,885 | 116,634 | 685,098 | 25,865 | 254,201 | 11,133 | 10,889 | 77.15% | 94.00% | 84.75% |
| | | | 30x | 414,336 | 96,183 | 726,328 | 25,784 | 273,730 | 10,860 | 12,025 | 81.16% | 94.30% | 87.24% |
| | | | 40x | 426,064 | 84,455 | 752,199 | 25,627 | 287,351 | 10,700 | 12,577 | 83.46% | 94.49% | 88.63% |
| | | | 50x | 433,887 | 76,632 | 769,851 | 25,551 | 296,823 | 10,539 | 12,874 | 84.99% | 94.60% | 89.54% |
| | HG005 | 416,777 | 10x | 318,752 | 98,025 | 572,191 | 22,606 | 223,966 | 12,069 | 6,153 | 76.48% | 93.51% | 84.14% |
| | | | 20x | 366,474 | 50,303 | 679,817 | 16,610 | 287,161 | 8,423 | 5,543 | 87.93% | 95.77% | 91.68% |
| | | | 30x | 380,766 | 36,011 | 725,133 | 13,644 | 320,165 | 6,872 | 4,948 | 91.36% | 96.63% | 93.92% |
| | | | 40x | 387,534 | 29,243 | 751,004 | 12,011 | 340,448 | 6,067 | 4,540 | 92.98% | 97.07% | 94.99% |
| | | | 50x | 391,699 | 25,078 | 767,723 | 11,023 | 353,719 | 5,552 | 4,253 | 93.98% | 97.34% | 95.63% |
| | HG006 | 428,237 | 10x | 306,570 | 121,668 | 576,385 | 22,399 | 240,721 | 11,029 | 6,378 | 71.59% | 93.33% | 81.03% |
| | | | 20x | 352,774 | 75,464 | 682,143 | 18,966 | 301,153 | 7,991 | 7,342 | 82.38% | 95.02% | 88.25% |
| | | | 30x | 368,436 | 59,802 | 728,163 | 18,120 | 331,388 | 7,234 | 7,940 | 86.04% | 95.43% | 90.49% |
| | | | 40x | 376,103 | 52,135 | 753,390 | 17,246 | 349,297 | 6,766 | 7,948 | 87.83% | 95.73% | 91.61% |
| | | | 50x | 381,562 | 46,676 | 770,120 | 16,726 | 360,816 | 6,488 | 7,938 | 89.10% | 95.91% | 92.38% |
| | HG007 | 430,326 | 10x | 306,268 | 124,058 | 578,077 | 22,589 | 242,380 | 11,025 | 6,592 | 71.17% | 93.27% | 80.74% |
| | | | 20x | 351,768 | 78,558 | 682,171 | 19,002 | 302,055 | 7,861 | 7,631 | 81.74% | 95.00% | 87.88% |
| | | | 30x | 366,308 | 64,018 | 722,554 | 17,879 | 328,174 | 7,224 | 8,017 | 85.12% | 95.47% | 90.00% |
| | | | 40x | 374,781 | 55,545 | 749,539 | 17,382 | 346,643 | 6,932 | 8,090 | 87.09% | 95.69% | 91.19% |
| | | | 50x | 380,556 | 49,770 | 768,128 | 17,239 | 359,342 | 6,775 | 8,258 | 88.43% | 95.78% | 91.96% |

***Supplementary table 4*** *Clair3 baseline (SUP) performance*

| Type | Sample | TRUTH.TOTAL | Depth | TRUTH.TP | TRUTH.FN | QUERY.TOTAL | QUERY.FP | QUERY.UNK | FP.gt | FP.al | METRIC.Recall | METRIC.Precision | METRIC.F1_Score |
|---|---|---|---|---|---|---|---|---|---|---|---|---|---|
| SNP | HG001 | 3,254,386 | 10x | 3,156,650 | 97,736 | 3,999,484 | 35,078 | 807,238 | 21,839 | 1,608 | 97.00% | 98.90% | 97.94% |
| | | | 20x | 3,233,130 | 21,256 | 4,170,266 | 11,154 | 925,175 | 4,566 | 699 | 99.35% | 99.66% | 99.50% |
| | | | 30x | 3,239,627 | 14,759 | 4,233,949 | 8,407 | 985,034 | 3,534 | 474 | 99.55% | 99.74% | 99.64% |
| | | | 40x | 3,242,817 | 11,569 | 4,275,352 | 7,975 | 1,023,634 | 3,319 | 429 | 99.64% | 99.75% | 99.70% |
| | | | 50x | 3,245,828 | 8,558 | 4,315,725 | 5,993 | 1,062,955 | 2,121 | 366 | 99.74% | 99.82% | 99.78% |
| | HG003 | 3,327,495 | 10x | 3,232,006 | 95,489 | 4,042,699 | 34,443 | 775,444 | 20,843 | 2,270 | 97.13% | 98.95% | 98.03% |
| | | | 20x | 3,297,116 | 30,379 | 4,203,264 | 11,487 | 893,507 | 5,037 | 1,074 | 99.09% | 99.65% | 99.37% |
| | | | 30x | 3,304,157 | 23,338 | 4,277,583 | 9,014 | 963,123 | 3,725 | 737 | 99.30% | 99.73% | 99.51% |
| | | | 40x | 3,308,906 | 18,589 | 4,319,473 | 7,821 | 1,001,360 | 3,276 | 585 | 99.44% | 99.76% | 99.60% |
| | | | 50x | 3,312,391 | 15,104 | 4,357,381 | 6,729 | 1,036,895 | 2,519 | 620 | 99.55% | 99.80% | 99.67% |
| | HG004 | 3,346,610 | 10x | 3,241,544 | 105,066 | 4,054,391 | 34,063 | 777,929 | 23,390 | 1,847 | 96.86% | 98.96% | 97.90% |
| | | | 20x | 3,315,381 | 31,229 | 4,227,149 | 11,193 | 899,346 | 5,355 | 909 | 99.07% | 99.66% | 99.36% |
| | | | 30x | 3,323,344 | 23,266 | 4,296,086 | 8,289 | 963,120 | 4,152 | 697 | 99.30% | 99.75% | 99.53% |
| | | | 40x | 3,329,047 | 17,563 | 4,346,758 | 6,698 | 1,009,651 | 3,559 | 617 | 99.48% | 99.80% | 99.64% |
| | | | 50x | 3,332,701 | 13,909 | 4,391,176 | 5,664 | 1,051,373 | 2,813 | 596 | 99.58% | 99.83% | 99.71% |
| | HG005 | 3,275,631 | 10x | 3,187,921 | 87,710 | 4,033,123 | 33,268 | 811,199 | 19,469 | 1,699 | 97.32% | 98.97% | 98.14% |
| | | | 20x | 3,251,995 | 23,636 | 4,180,266 | 10,570 | 916,666 | 4,951 | 707 | 99.28% | 99.68% | 99.48% |
| | | | 30x | 3,258,477 | 17,154 | 4,252,324 | 8,750 | 983,986 | 3,977 | 619 | 99.48% | 99.73% | 99.60% |
| | | | 40x | 3,262,329 | 13,302 | 4,305,768 | 7,620 | 1,034,676 | 3,386 | 540 | 99.59% | 99.77% | 99.68% |
| | | | 50x | 3,265,054 | 10,577 | 4,348,935 | 6,572 | 1,076,176 | 2,691 | 494 | 99.68% | 99.80% | 99.74% |
| | HG006 | 3,269,860 | 10x | 3,179,957 | 89,903 | 3,992,292 | 35,280 | 776,637 | 20,305 | 1,651 | 97.25% | 98.90% | 98.07% |
| | | | 20x | 3,243,847 | 26,013 | 4,137,678 | 11,556 | 881,696 | 5,079 | 665 | 99.20% | 99.65% | 99.42% |
| | | | 30x | 3,250,032 | 19,828 | 4,219,484 | 9,030 | 959,773 | 4,149 | 512 | 99.39% | 99.72% | 99.56% |
| | | | 40x | 3,254,598 | 15,262 | 4,274,385 | 8,683 | 1,010,428 | 3,601 | 529 | 99.53% | 99.73% | 99.63% |
| | | | 50x | 3,257,855 | 12,005 | 4,311,895 | 7,026 | 1,046,326 | 2,766 | 450 | 99.63% | 99.78% | 99.71% |
| | HG007 | 3,284,462 | 10x | 3,190,577 | 93,885 | 4,004,241 | 32,935 | 780,391 | 21,172 | 1,437 | 97.14% | 98.98% | 98.05% |
| | | | 20x | 3,254,713 | 29,749 | 4,147,757 | 10,203 | 882,344 | 4,697 | 652 | 99.09% | 99.69% | 99.39% |
| | | | 30x | 3,261,346 | 23,116 | 4,207,326 | 8,328 | 937,092 | 4,070 | 493 | 99.30% | 99.75% | 99.52% |
| | | | 40x | 3,266,605 | 17,857 | 4,269,809 | 7,211 | 995,391 | 3,436 | 435 | 99.46% | 99.78% | 99.62% |
| | | | 50x | 3,270,349 | 14,113 | 4,312,548 | 5,995 | 1,035,585 | 2,643 | 408 | 99.57% | 99.82% | 99.69% |
| INDEL | HG001 | 467,702 | 10x | 302,763 | 164,939 | 495,245 | 15,430 | 172,586 | 6,727 | 3,912 | 64.73% | 95.22% | 77.07% |
| | | | 20x | 358,114 | 109,588 | 597,014 | 11,629 | 220,389 | 4,645 | 4,156 | 76.57% | 96.91% | 85.55% |
| | | | 30x | 376,752 | 90,950 | 637,358 | 10,983 | 241,611 | 4,158 | 4,606 | 80.55% | 97.22% | 88.11% |
| | | | 40x | 387,302 | 80,400 | 660,953 | 10,572 | 254,347 | 3,943 | 4,843 | 82.81% | 97.40% | 89.51% |
| | | | 50x | 394,396 | 73,306 | 677,505 | 10,500 | 263,519 | 3,798 | 5,059 | 84.33% | 97.46% | 90.42% |
| | HG003 | 504,501 | 10x | 324,992 | 179,509 | 514,406 | 16,829 | 166,776 | 7,256 | 4,557 | 64.42% | 95.16% | 76.83% |
| | | | 20x | 379,958 | 124,543 | 607,678 | 12,595 | 206,689 | 5,028 | 4,764 | 75.31% | 96.86% | 84.74% |
| | | | 30x | 399,586 | 104,915 | 647,007 | 12,088 | 225,575 | 4,709 | 5,272 | 79.20% | 97.13% | 87.26% |
| | | | 40x | 410,591 | 93,910 | 669,463 | 11,771 | 236,732 | 4,455 | 5,605 | 81.39% | 97.28% | 88.63% |
| | | | 50x | 418,095 | 86,406 | 685,023 | 11,954 | 244,231 | 4,448 | 5,896 | 82.87% | 97.29% | 89.50% |
| | HG004 | 510,519 | 10x | 323,802 | 186,717 | 511,068 | 17,094 | 164,506 | 7,567 | 4,515 | 63.43% | 95.07% | 76.09% |
| | | | 20x | 380,421 | 130,098 | 607,883 | 12,840 | 206,103 | 5,076 | 4,794 | 74.52% | 96.80% | 84.21% |
| | | | 30x | 399,860 | 110,659 | 645,048 | 12,281 | 223,166 | 4,604 | 5,473 | 78.32% | 97.09% | 86.70% |
| | | | 40x | 411,521 | 98,998 | 668,949 | 12,232 | 234,684 | 4,513 | 5,886 | 80.61% | 97.18% | 88.12% |
| | | | 50x | 419,354 | 91,165 | 686,040 | 12,396 | 243,405 | 4,414 | 6,250 | 82.14% | 97.20% | 89.04% |
| | HG005 | 416,777 | 10x | 309,324 | 107,453 | 506,588 | 14,134 | 178,100 | 6,247 | 3,306 | 74.22% | 95.70% | 83.60% |
| | | | 20x | 357,966 | 58,811 | 601,594 | 8,804 | 227,400 | 3,706 | 2,823 | 85.89% | 97.65% | 91.39% |
| | | | 30x | 372,667 | 44,110 | 641,528 | 7,200 | 253,183 | 2,896 | 2,686 | 89.42% | 98.15% | 93.58% |
| | | | 40x | 379,413 | 37,364 | 663,551 | 6,166 | 268,990 | 2,467 | 2,471 | 91.04% | 98.44% | 94.59% |
| | | | 50x | 383,528 | 33,249 | 678,400 | 5,757 | 279,900 | 2,265 | 2,434 | 92.02% | 98.56% | 95.18% |
| | HG006 | 428,237 | 10x | 297,865 | 130,373 | 509,448 | 13,932 | 192,606 | 5,789 | 3,414 | 69.56% | 95.60% | 80.53% |
| | | | 20x | 343,746 | 84,492 | 603,389 | 9,359 | 243,001 | 3,493 | 3,344 | 80.27% | 97.40% | 88.01% |
| | | | 30x | 358,883 | 69,355 | 642,506 | 8,224 | 267,193 | 2,882 | 3,520 | 83.80% | 97.81% | 90.27% |
| | | | 40x | 366,912 | 61,326 | 664,307 | 7,675 | 281,106 | 2,611 | 3,634 | 85.68% | 98.00% | 91.43% |
| | | | 50x | 372,738 | 55,500 | 679,886 | 7,461 | 290,817 | 2,499 | 3,734 | 87.04% | 98.08% | 92.23% |
| | HG007 | 430,326 | 10x | 297,049 | 133,277 | 510,791 | 14,215 | 194,492 | 5,890 | 3,603 | 69.03% | 95.51% | 80.14% |
| | | | 20x | 342,616 | 87,710 | 604,147 | 9,700 | 244,446 | 3,514 | 3,608 | 79.62% | 97.30% | 87.58% |
| | | | 30x | 356,825 | 73,501 | 639,556 | 8,415 | 266,078 | 3,054 | 3,607 | 82.92% | 97.75% | 89.72% |
| | | | 40x | 365,421 | 64,905 | 663,276 | 8,223 | 280,928 | 2,848 | 3,887 | 84.92% | 97.85% | 90.93% |
| | | | 50x | 371,478 | 58,848 | 679,978 | 8,120 | 291,425 | 2,702 | 4,053 | 86.32% | 97.91% | 91.75% |

***Supplementary table 5*** *Clair3 (HAC) indel performance comparison stratified by length*

| | | F1 score | | Recall | | Precision | |
|---|---|---|---|---|---|---|---|
| **Depth** | **Category** | **With dwelling time** | **Without dwelling time** | **With dwelling time** | **Without dwelling time** | **With dwelling time** | **Without dwelling time** |
| | SNP | 97.78% | 96.40% | 96.80% | 94.36% | 98.78% | 98.53% |
| | Del 1-5bp | 80.34% | 67.12% | 70.73% | 51.18% | 92.98% | 97.47% |
| | Del 6-15bp | 85.47% | 68.81% | 80.95% | 53.67% | 90.53% | 95.86% |
| | Del 16+bp | 88.84% | 70.36% | 83.14% | 55.02% | 95.39% | 97.56% |
| | Ins 1-5bp | 79.86% | 68.28% | 70.62% | 53.07% | 91.88% | 95.72% |
| | Ins 6-15bp | 82.04% | 54.88% | 79.18% | 39.09% | 85.10% | 92.08% |
| | Ins 16+bp | 87.46% | 63.46% | 83.79% | 47.65% | 91.47% | 94.95% |
| 10x | All INDELs | 81.09% | 68.04% | 72.40% | 52.56% | 92.15% | 96.45% |
| | SNP | 99.51% | 99.20% | 99.28% | 98.83% | 99.74% | 99.57% |
| | Del 1-5bp | 88.99% | 80.92% | 83.31% | 68.63% | 95.50% | 98.58% |
| | Del 6-15bp | 93.51% | 84.98% | 92.02% | 75.66% | 95.05% | 96.92% |
| | Del 16+bp | 95.63% | 86.60% | 93.67% | 77.61% | 97.67% | 97.93% |
| | Ins 1-5bp | 88.32% | 80.34% | 82.88% | 68.46% | 94.53% | 97.21% |
| | Ins 6-15bp | 91.46% | 74.72% | 91.25% | 62.70% | 91.67% | 92.44% |
| | Ins 16+bp | 94.52% | 83.05% | 93.22% | 72.86% | 95.85% | 96.56% |
| 20x | All INDELs | 89.43% | 81.36% | 84.46% | 69.72% | 95.02% | 97.68% |
| | SNP | 99.67% | 99.52% | 99.53% | 99.37% | 99.81% | 99.67% |
| | Del 1-5bp | 91.64% | 85.43% | 87.49% | 75.21% | 96.20% | 98.86% |
| | Del 6-15bp | 95.74% | 89.87% | 94.91% | 83.40% | 96.59% | 97.42% |
| | Del 16+bp | 97.23% | 90.36% | 96.11% | 83.88% | 98.39% | 97.92% |
| | Ins 1-5bp | 91.10% | 83.80% | 87.21% | 73.42% | 95.37% | 97.61% |
| | Ins 6-15bp | 94.23% | 80.99% | 94.40% | 72.02% | 94.06% | 92.50% |
| | Ins 16+bp | 96.28% | 87.61% | 95.55% | 80.00% | 97.03% | 96.81% |
| 30x | All INDELs | 92.01% | 85.35% | 88.44% | 75.59% | 95.88% | 98.01% |
| | SNP | 99.74% | 99.61% | 99.65% | 99.51% | 99.83% | 99.70% |
| | Del 1-5bp | 92.97% | 87.53% | 89.63% | 78.49% | 96.56% | 98.92% |
| | Del 6-15bp | 96.78% | 91.96% | 96.21% | 86.80% | 97.35% | 97.78% |
| | Del 16+bp | 97.89% | 91.29% | 97.15% | 85.84% | 98.64% | 97.49% |
| | Ins 1-5bp | 92.46% | 85.41% | 89.33% | 75.86% | 95.82% | 97.70% |
| | Ins 6-15bp | 95.57% | 83.52% | 95.76% | 76.36% | 95.38% | 92.16% |
| | Ins 16+bp | 97.33% | 89.46% | 96.83% | 83.05% | 97.83% | 96.94% |
| 40x | All INDELs | 93.29% | 87.17% | 90.43% | 78.44% | 96.33% | 98.09% |
| | SNP | 99.81% | 99.65% | 99.75% | 99.56% | 99.86% | 99.73% |
| | Del 1-5bp | 93.80% | 88.66% | 90.95% | 80.28% | 96.83% | 99.00% |
| | Del 6-15bp | 97.34% | 92.82% | 96.81% | 88.17% | 97.88% | 98.00% |
| | Del 16+bp | 98.09% | 91.88% | 97.39% | 86.64% | 98.80% | 97.80% |
| | Ins 1-5bp | 93.21% | 86.21% | 90.47% | 77.07% | 96.12% | 97.81% |
| | Ins 6-15bp | 96.04% | 84.39% | 96.22% | 78.00% | 95.86% | 91.92% |
| | Ins 16+bp | 97.54% | 89.99% | 97.01% | 83.92% | 98.06% | 97.02% |
| 50x | All INDELs | 94.02% | 88.10% | 91.54% | 79.89% | 96.63% | 98.18% |

***Supplementary table 6*** *Clair3 (SUP) indel performance comparison stratified by length*

| Depth | Category | F1 score | | Recall | | Precision | |
|---|---|---|---|---|---|---|---|
| | | With dwelling time | Without dwelling time | With dwelling time | Without dwelling time | With dwelling time | Without dwelling time |
| 10x | SNP | 98.34% | 98.14% | 97.66% | 97.32% | 99.02% | 98.97% |
| | Del 1-5bp | 83.83% | 82.70% | 75.73% | 72.29% | 93.86% | 96.61% |
| | Del 6-15bp | 87.67% | 81.46% | 84.37% | 70.47% | 91.24% | 96.52% |
| | Del 16+bp | 92.22% | 83.22% | 88.52% | 72.45% | 96.25% | 97.74% |
| | Ins 1-5bp | 82.83% | 83.59% | 74.05% | 74.80% | 93.98% | 94.73% |
| | Ins 6-15bp | 82.81% | 81.38% | 79.52% | 71.63% | 86.38% | 94.20% |
| | Ins 16+bp | 89.99% | 88.19% | 86.82% | 81.78% | 93.40% | 95.71% |
| | All INDELs | 84.14% | 83.60% | 76.48% | 74.22% | 93.51% | 95.70% |
| 20x | SNP | 99.68% | 99.48% | 99.57% | 99.28% | 99.78% | 99.68% |
| | Del 1-5bp | 91.48% | 90.78% | 87.57% | 84.40% | 95.75% | 98.21% |
| | Del 6-15bp | 95.08% | 91.70% | 94.30% | 85.74% | 95.87% | 98.56% |
| | Del 16+bp | 97.40% | 94.48% | 96.65% | 90.75% | 98.16% | 98.53% |
| | Ins 1-5bp | 90.68% | 91.06% | 85.98% | 85.84% | 95.93% | 96.96% |
| | Ins 6-15bp | 91.76% | 90.76% | 91.52% | 85.01% | 92.00% | 97.34% |
| | Ins 16+bp | 96.22% | 94.04% | 95.55% | 90.86% | 96.91% | 97.46% |
| | All INDELs | 91.68% | 91.39% | 87.93% | 85.89% | 95.77% | 97.65% |
| 30x | SNP | 99.79% | 99.60% | 99.74% | 99.48% | 99.84% | 99.73% |
| | Del 1-5bp | 93.77% | 93.16% | 91.25% | 88.28% | 96.43% | 98.61% |
| | Del 6-15bp | 96.98% | 94.29% | 96.57% | 89.95% | 97.39% | 99.06% |
| | Del 16+bp | 98.16% | 96.24% | 97.75% | 93.91% | 98.58% | 98.69% |
| | Ins 1-5bp | 93.08% | 93.22% | 89.67% | 89.28% | 96.77% | 97.53% |
| | Ins 6-15bp | 94.42% | 93.20% | 94.39% | 88.65% | 94.45% | 98.24% |
| | Ins 16+bp | 97.65% | 95.08% | 97.28% | 92.38% | 98.03% | 97.94% |
| | All INDELs | 93.92% | 93.58% | 91.36% | 89.42% | 96.63% | 98.15% |
| 40x | SNP | 99.83% | 99.68% | 99.81% | 99.59% | 99.86% | 99.77% |
| | Del 1-5bp | 94.78% | 94.28% | 92.87% | 90.15% | 96.77% | 98.82% |
| | Del 6-15bp | 97.87% | 95.19% | 97.50% | 91.46% | 98.25% | 99.23% |
| | Del 16+bp | 98.43% | 96.30% | 98.27% | 94.04% | 98.59% | 98.67% |
| | Ins 1-5bp | 94.32% | 94.27% | 91.58% | 90.88% | 97.22% | 97.93% |
| | Ins 6-15bp | 95.65% | 94.25% | 95.67% | 90.30% | 95.63% | 98.56% |
| | Ins 16+bp | 98.09% | 95.55% | 97.67% | 92.96% | 98.50% | 98.30% |
| | All INDELs | 94.99% | 94.59% | 92.98% | 91.04% | 97.07% | 98.44% |
| 50x | SNP | 99.86% | 99.74% | 99.84% | 99.68% | 99.88% | 99.80% |
| | Del 1-5bp | 95.46% | 94.94% | 93.94% | 91.27% | 97.03% | 98.92% |
| | Del 6-15bp | 98.38% | 95.55% | 98.06% | 91.95% | 98.70% | 99.45% |
| | Del 16+bp | 98.64% | 96.24% | 98.46% | 93.84% | 98.83% | 98.77% |
| | Ins 1-5bp | 94.99% | 94.87% | 92.67% | 91.90% | 97.43% | 98.05% |
| | Ins 6-15bp | 96.30% | 94.79% | 96.33% | 91.16% | 96.28% | 98.72% |
| | Ins 16+bp | 98.28% | 95.81% | 97.98% | 93.39% | 98.59% | 98.35% |
| | All INDELs | 95.63% | 95.18% | 93.98% | 92.02% | 97.34% | 98.56% |

***Supplementary table 7*** *Clair3 v2 performance in complex regions (HG005)*

| Variant type | Enable dwellign time? | Stratification type | Region | TP | FP | FN | Precision | Recall | F1 |
|---|---|---|---|---|---|---|---|---|---|
| | | | Homopol_4-6bp | 112,166 | 2,970 | 8,143 | 97.46% | 93.23% | 95.30% |
| | | | Homopol_7-11bp | 79,649 | 5,123 | 13,238 | 93.99% | 85.75% | 89.68% |
| | | | Homopol_ge12bp | 14,356 | 6,148 | 28,730 | 70.39% | 33.32% | 45.23% |
| | | | Imp_Homopol_ge11bp | 40,806 | 8,643 | 35,721 | 82.77% | 53.32% | 64.86% |
| | | | TR_le50bp | 57,080 | 3,084 | 3,496 | 95.35% | 94.23% | 94.79% |
| | | | TR51-200bp | 25,693 | 1,348 | 1,814 | 95.46% | 93.41% | 94.42% |
| | | | TR201-10kbp | 3,981 | 49 | 100 | 98.82% | 97.55% | 98.18% |
| | | | TR_ge101bp | 11,674 | 299 | 516 | 97.63% | 95.77% | 96.69% |
| | | LowComplexity | TR_and_Homopol | 186,577 | 15,712 | 46,968 | 92.59% | 79.89% | 85.77% |
| | | | SegDups | 9,962 | 359 | 399 | 96.57% | 96.15% | 96.36% |
| | | SegmentalDuplications | SegDups_gt10kb | 8,472 | 332 | 358 | 96.28% | 95.95% | 96.11% |
| | | Mappability | LowMap | 9,500 | 217 | 299 | 97.77% | 96.95% | 97.36% |
| | | | L1H | 196 | 1 | 3 | 99.49% | 98.49% | 98.99% |
| | | OtherDifficult | MHC | 1,552 | 72 | 146 | 95.87% | 91.40% | 93.58% |
| | Yes | FunctionalRegions | CDS | 352 | 5 | 5 | 98.63% | 98.60% | 98.61% |
| | | | Homopol_4-6bp | 100,733 | 1,433 | 19,573 | 98.61% | 83.73% | 90.56% |
| | | | Homopol_7-11bp | 58,468 | 1,727 | 34,418 | 97.13% | 62.95% | 76.39% |
| | | | Homopol_ge12bp | 3,374 | 763 | 39,712 | 81.49% | 7.83% | 14.29% |
| | | | Imp_Homopol_ge11bp | 20,628 | 1,520 | 55,899 | 93.17% | 26.96% | 41.81% |
| | | | TR_le50bp | 45,919 | 2,357 | 14,657 | 95.51% | 75.80% | 84.52% |
| | | | TR51-200bp | 20,707 | 824 | 6,800 | 96.46% | 75.28% | 84.56% |
| | | | TR201-10kbp | 3,632 | 57 | 449 | 98.49% | 89.00% | 93.51% |
| | | | TR_ge101bp | 10,063 | 232 | 2,127 | 97.84% | 82.55% | 89.55% |
| | | LowComplexity | TR_and_Homopol | 137,413 | 5,809 | 96,132 | 96.11% | 58.84% | 72.99% |
| | | | SegDups | 9,230 | 279 | 1,131 | 97.10% | 89.08% | 92.92% |
| | | SegmentalDuplications | SegDups_gt10kb | 7,852 | 259 | 978 | 96.84% | 88.92% | 92.72% |
| | | Mappability | LowMap | 8,990 | 188 | 809 | 97.95% | 91.74% | 94.75% |
| | | | L1H | 186 | 1 | 13 | 99.47% | 93.47% | 96.37% |
| | | OtherDifficult | MHC | 1,389 | 57 | 309 | 96.30% | 81.80% | 88.46% |
| Indel | No | FunctionalRegions | CDS | 324 | 4 | 33 | 98.78% | 90.76% | 94.60% |
| | | | Homopol_4-6bp | 812,358 | 1,623 | 3,729 | 99.80% | 99.54% | 99.67% |
| | | | Homopol_7-11bp | 62,894 | 547 | 794 | 99.14% | 98.75% | 98.95% |
| | | | Homopol_ge12bp | 5,064 | 434 | 751 | 92.27% | 87.09% | 89.60% |
| | | | Imp_Homopol_ge11bp | 27,187 | 650 | 1,091 | 97.68% | 96.14% | 96.91% |
| | | | TR_le50bp | 30,787 | 131 | 283 | 99.58% | 99.09% | 99.33% |
| | | | TR51-200bp | 24,782 | 124 | 254 | 99.51% | 98.99% | 99.25% |
| | | | TR201-10kbp | 10,212 | 36 | 39 | 99.65% | 99.62% | 99.63% |
| | | | TR_ge101bp | 19,702 | 66 | 136 | 99.67% | 99.31% | 99.49% |
| | | LowComplexity | TR_and_Homopol | 146,118 | 1,224 | 2,118 | 99.18% | 98.57% | 98.87% |
| | | | SegDups | 121,534 | 3,319 | 2,581 | 97.34% | 97.92% | 97.63% |
| | | SegmentalDuplications | SegDups_gt10kb | 104,972 | 3,301 | 2,535 | 96.95% | 97.64% | 97.30% |
| | | Mappability | LowMap | 190,184 | 2,416 | 2,778 | 98.75% | 98.56% | 98.65% |
| | | | L1H | 5,574 | 5 | 29 | 99.83% | 99.48% | 99.70% |
| | | OtherDifficult | MHC | 18,685 | 43 | 200 | 99.77% | 98.94% | 99.35% |
| | Yes | FunctionalRegions | CDS | 20,536 | 35 | 129 | 99.83% | 99.38% | 99.60% |
| | | | Homopol_4-6bp | 809,263 | 3,268 | 6,824 | 99.60% | 99.16% | 99.38% |
| | | | Homopol_7-11bp | 59,794 | 1,760 | 3,894 | 97.15% | 93.89% | 95.49% |
| | | | Homopol_ge12bp | 3,856 | 875 | 1,959 | 81.69% | 66.31% | 73.20% |
| | | | Imp_Homopol_ge11bp | 24,874 | 1,498 | 3,404 | 94.34% | 87.96% | 91.04% |
| | | | TR_le50bp | 29,968 | 447 | 1,102 | 98.53% | 96.45% | 97.48% |
| | | | TR51-200bp | 24,251 | 360 | 785 | 98.54% | 96.86% | 97.70% |
| | | | TR201-10kbp | 10,165 | 68 | 86 | 99.33% | 99.16% | 99.25% |
| | | | TR_ge101bp | 19,493 | 178 | 345 | 99.10% | 98.26% | 98.68% |
| | | LowComplexity | TR_and_Homopol | 140,361 | 3,477 | 7,875 | 97.59% | 94.69% | 96.12% |
| | | | SegDups | 121,068 | 4,053 | 3,047 | 96.76% | 97.55% | 97.15% |
| | | SegmentalDuplications | SegDups_gt10kb | 104,523 | 3,996 | 2,984 | 96.32% | 97.22% | 96.77% |
| | | Mappability | LowMap | 189,805 | 2,904 | 3,157 | 98.49% | 98.36% | 98.43% |
| | | | L1H | 5,573 | 8 | 30 | 99.86% | 99.46% | 99.66% |
| | | OtherDifficult | MHC | 18,530 | 94 | 355 | 99.49% | 98.12% | 98.80% |
| SNP | No | FunctionalRegions | CDS | 20,575 | 61 | 90 | 99.70% | 99.56% | 99.63% |

***Supplementary Table 8*** *Clair3 v2 performance in complex regions (HG001)*

| Variant type | Enable dwellign time? | Stratification type | Region | TP | FP | FN | Precision | Recall | F1 |
|---|---|---|---|---|---|---|---|---|---|
| Indel | Yes | LowComplexity | Homopol_4-6bp | 111,116 | 3,913 | 14,950 | 96.64% | 88.14% | 92.20% |
| | | | Homopol_7-11bp | 82,568 | 5,910 | 16,443 | 93.36% | 83.39% | 88.09% |
| | | | Homopol_ge12bp | 21,954 | 13,211 | 73,730 | 62.88% | 22.94% | 33.62% |
| | | | Imp_Homopol_ge11bp | 49,687 | 16,253 | 82,399 | 75.64% | 37.62% | 50.25% |
| | | | TR_le50bp | 54,023 | 3,074 | 3,497 | 95.09% | 93.92% | 94.50% |
| | | | TR51-200bp | 25,400 | 1,483 | 2,120 | 94.97% | 92.30% | 93.61% |
| | | | TR201-10kbp | 4,141 | 61 | 114 | 98.59% | 97.32% | 97.95% |
| | | | TR_ge101bp | 12,248 | 403 | 649 | 96.98% | 94.97% | 95.96% |
| | | | TR_and_Homopol | 193,172 | 23,557 | 94,343 | 89.58% | 67.19% | 76.78% |
| | | SegmentalDuplications | SegDups | 8,850 | 380 | 290 | 95.94% | 96.83% | 96.38% |
| | | | SegDups_gt10kb | 7,541 | 364 | 259 | 95.47% | 96.68% | 96.07% |
| | | Mappability | LowMap | 8,655 | 176 | 218 | 98.02% | 97.54% | 97.78% |
| | | OtherDifficult | L1H | 196 | 2 | 2 | 98.99% | 98.99% | 98.99% |
| | | | MHC | 1,693 | 89 | 139 | 95.26% | 92.41% | 93.82% |
| | | FunctionalRegions | CDS | 351 | 10 | 5 | 97.22% | 98.60% | 97.90% |
| | No | LowComplexity | Homopol_4-6bp | 98,436 | 1,500 | 27,623 | 98.51% | 78.09% | 87.12% |
| | | | Homopol_7-11bp | 59,785 | 1,846 | 39,225 | 97.01% | 60.38% | 74.43% |
| | | | Homopol_ge12bp | 4,191 | 1,302 | 91,493 | 76.21% | 4.38% | 8.28% |
| | | | Imp_Homopol_ge11bp | 21,883 | 2,132 | 110,203 | 91.14% | 16.57% | 28.04% |
| | | | TR_le50bp | 43,066 | 2,265 | 14,454 | 95.39% | 74.87% | 83.90% |
| | | | TR51-200bp | 19,977 | 878 | 7,543 | 96.10% | 72.59% | 82.71% |
| | | | TR201-10kbp | 3,762 | 65 | 493 | 98.34% | 88.41% | 93.12% |
| | | | TR_ge101bp | 10,345 | 310 | 2,552 | 97.21% | 80.21% | 87.90% |
| | | | TR_and_Homopol | 135,682 | 6,398 | 151,833 | 95.67% | 47.19% | 63.21% |
| | | SegmentalDuplications | SegDups | 8,124 | 333 | 1,016 | 96.11% | 88.88% | 92.36% |
| | | | SegDups_gt10kb | 6,935 | 321 | 865 | 95.63% | 88.91% | 92.15% |
| | | Mappability | LowMap | 8,223 | 160 | 650 | 98.10% | 92.67% | 95.31% |
| | | OtherDifficult | L1H | 189 | 2 | 9 | 98.95% | 95.45% | 97.17% |
| | | | MHC | 1,497 | 51 | 335 | 96.86% | 81.71% | 88.64% |
| | | FunctionalRegions | CDS | 321 | 4 | 35 | 98.76% | 90.17% | 94.27% |
| SNP | Yes | LowComplexity | Homopol_4-6bp | 804,723 | 1,619 | 3,418 | 99.80% | 99.58% | 99.69% |
| | | | Homopol_7-11bp | 62,327 | 623 | 891 | 99.02% | 98.59% | 98.80% |
| | | | Homopol_ge12bp | 8,843 | 646 | 1,665 | 93.29% | 84.15% | 88.49% |
| | | | Imp_Homopol_ge11bp | 31,932 | 942 | 2,087 | 97.16% | 93.87% | 95.48% |
| | | | TR_le50bp | 30,313 | 122 | 280 | 99.60% | 99.08% | 99.34% |
| | | | TR51-200bp | 25,580 | 147 | 298 | 99.43% | 98.85% | 99.14% |
| | | | TR201-10kbp | 10,353 | 24 | 63 | 99.77% | 99.40% | 99.58% |
| | | | TR_ge101bp | 20,292 | 71 | 167 | 99.65% | 99.18% | 99.42% |
| | | | TR_and_Homopol | 149,819 | 1,517 | 3,159 | 99.00% | 97.94% | 98.47% |
| | | SegmentalDuplications | SegDups | 110,192 | 3,623 | 614 | 96.82% | 99.45% | 98.11% |
| | | | SegDups_gt10kb | 94,074 | 3,594 | 566 | 96.32% | 99.40% | 97.84% |
| | | Mappability | LowMap | 174,871 | 1,931 | 851 | 98.91% | 99.52% | 99.21% |
| | | OtherDifficult | L1H | 5,454 | 7 | 28 | 99.87% | 99.49% | 99.68% |
| | | | MHC | 20,283 | 37 | 167 | 99.82% | 99.18% | 99.50% |
| | | FunctionalRegions | CDS | 20,098 | 37 | 95 | 99.82% | 99.53% | 99.67% |
| | No | LowComplexity | Homopol_4-6bp | 801,460 | 2,877 | 6,676 | 99.64% | 99.17% | 99.41% |
| | | | Homopol_7-11bp | 59,240 | 1,662 | 3,979 | 97.28% | 93.71% | 95.46% |
| | | | Homopol_ge12bp | 7,002 | 1,033 | 3,506 | 87.25% | 66.63% | 75.56% |
| | | | Imp_Homopol_ge11bp | 28,847 | 1,673 | 5,172 | 94.54% | 84.80% | 89.40% |
| | | | TR_le50bp | 29,577 | 328 | 1,016 | 98.90% | 96.68% | 97.78% |
| | | | TR51-200bp | 25,027 | 359 | 851 | 98.59% | 96.71% | 97.64% |
| | | | TR201-10kbp | 10,317 | 42 | 99 | 99.59% | 99.05% | 99.32% |
| | | | TR_ge101bp | 20,073 | 156 | 386 | 99.23% | 98.11% | 98.67% |
| | | | TR_and_Homopol | 143,504 | 3,380 | 9,474 | 97.70% | 93.81% | 95.72% |
| | | SegmentalDuplications | SegDups | 110,165 | 3,557 | 641 | 96.87% | 99.42% | 98.13% |
| | | | SegDups_gt10kb | 94,048 | 3,521 | 592 | 96.39% | 99.37% | 97.86% |
| | | Mappability | LowMap | 174,883 | 1,935 | 839 | 98.91% | 99.52% | 99.21% |
| | | OtherDifficult | L1H | 5,461 | 14 | 21 | 99.74% | 99.62% | 99.68% |
| | | | MHC | 20,131 | 85 | 319 | 99.58% | 98.44% | 99.01% |
| | | FunctionalRegions | CDS | 20,111 | 54 | 82 | 99.73% | 99.59% | 99.66% |

***Supplementary table 9*** *Different variant caller comparison*

| Varaint type | Depth | Variant caller | Sample | Region | TP | FP | FN | Precision | Recall | F1 score |
|---|---|---|---|---|---|---|---|---|---|---|
| | | Clair3 V2 | HG003 | chr20 | 68,753 | 822 | 1,413 | 98.82% | 97.99% | 98.40% |
| | | Dorado Variant | HG003 | chr20 | 65,266 | 3,306 | 4,900 | 95.10% | 93.02% | 94.05% |
| | | DeepVariant | HG003 | chr20 | 67,233 | 656 | 2,933 | 99.03% | 95.82% | 97.40% |
| | 10x | LongcallD | HG003 | chr20 | 64,198 | 2,069 | 5,968 | 96.88% | 91.49% | 94.11% |
| | | Clair3 V2 | HG003 | chr20 | 69,875 | 154 | 291 | 99.78% | 99.59% | 99.68% |
| | | Dorado Variant | HG003 | chr20 | 69,879 | 216 | 287 | 99.69% | 99.59% | 99.64% |
| | | DeepVariant | HG003 | chr20 | 69,994 | 137 | 172 | 99.80% | 99.75% | 99.78% |
| | 20x | LongcallD | HG003 | chr20 | 69,589 | 448 | 577 | 99.36% | 99.18% | 99.27% |
| | | Clair3 V2 | HG003 | chr20 | 69,981 | 133 | 185 | 99.81% | 99.74% | 99.77% |
| | | Dorado Variant | HG003 | chr20 | 70,015 | 104 | 151 | 99.85% | 99.78% | 99.82% |
| | | DeepVariant | HG003 | chr20 | 70,070 | 129 | 96 | 99.82% | 99.86% | 99.84% |
| | 30x | LongcallD | HG003 | chr20 | 69,916 | 281 | 250 | 99.60% | 99.64% | 99.62% |
| | | Clair3 V2 | HG003 | chr20 | 69,990 | 128 | 176 | 99.82% | 99.75% | 99.78% |
| | | Dorado Variant | HG003 | chr20 | 70,038 | 121 | 128 | 99.82% | 99.82% | 99.82% |
| | | DeepVariant | HG003 | chr20 | 70,075 | 123 | 91 | 99.82% | 99.87% | 99.85% |
| | 40x | LongcallD | HG003 | chr20 | 69,973 | 246 | 193 | 99.65% | 99.72% | 99.69% |
| | | Clair3 V2 | HG003 | chr20 | 70,025 | 121 | 141 | 99.83% | 99.80% | 99.81% |
| | | Dorado Variant | HG003 | chr20 | 70,026 | 132 | 140 | 99.81% | 99.80% | 99.80% |
| | | DeepVariant | HG003 | chr20 | 70,077 | 86 | 89 | 99.88% | 99.87% | 99.88% |
| SNP | 50x | LongcallD | HG003 | chr20 | 70,007 | 225 | 159 | 99.68% | 99.77% | 99.73% |
| | | Clair3 V2 | HG003 | chr20 | 6,696 | 547 | 3,932 | 92.44% | 63.00% | 74.93% |
| | | Dorado Variant | HG003 | chr20 | 5,535 | 3,502 | 5,093 | 61.23% | 52.08% | 56.29% |
| | | DeepVariant | HG003 | chr20 | 6,782 | 1,371 | 3,846 | 83.17% | 63.81% | 72.22% |
| | 10x | LongcallD | HG003 | chr20 | 6,729 | 10,749 | 3,899 | 39.10% | 63.31% | 48.34% |
| | | Clair3 V2 | HG003 | chr20 | 7,753 | 347 | 2,875 | 95.71% | 72.95% | 82.79% |
| | | Dorado Variant | HG003 | chr20 | 7,924 | 1,560 | 2,704 | 83.54% | 74.56% | 78.79% |
| | | DeepVariant | HG003 | chr20 | 8,053 | 1,510 | 2,575 | 84.20% | 75.77% | 79.76% |
| | 20x | LongcallD | HG003 | chr20 | 7,840 | 7,088 | 2,788 | 53.22% | 73.77% | 61.83% |
| | | Clair3 V2 | HG003 | chr20 | 8,249 | 332 | 2,379 | 96.13% | 77.62% | 85.89% |
| | | Dorado Variant | HG003 | chr20 | 8,542 | 1,388 | 2,086 | 86.01% | 80.37% | 83.09% |
| | | DeepVariant | HG003 | chr20 | 8,512 | 1,491 | 2,116 | 85.09% | 80.09% | 82.51% |
| | 30x | LongcallD | HG003 | chr20 | 8,100 | 5,900 | 2,528 | 58.58% | 76.21% | 66.24% |
| | | Clair3 V2 | HG003 | chr20 | 8,516 | 382 | 2,112 | 95.70% | 80.13% | 87.23% |
| | | Dorado Variant | HG003 | chr20 | 8,861 | 1,212 | 1,767 | 87.96% | 83.37% | 85.60% |
| | | DeepVariant | HG003 | chr20 | 8,780 | 1,450 | 1,848 | 85.82% | 82.61% | 84.18% |
| | 40x | LongcallD | HG003 | chr20 | 8,263 | 5,457 | 2,365 | 60.96% | 77.75% | 68.34% |
| | | Clair3 V2 | HG003 | chr20 | 8,786 | 390 | 1,842 | 95.75% | 82.67% | 88.73% |
| | | Dorado Variant | HG003 | chr20 | 8,968 | 1,148 | 1,660 | 88.64% | 84.38% | 86.46% |
| | | DeepVariant | HG003 | chr20 | 8,917 | 1,443 | 1,711 | 86.06% | 83.90% | 84.97% |
| Indel | 50x | LongcallD | HG003 | chr20 | 8,335 | 5,092 | 2,293 | 62.83% | 78.42% | 69.77% |

### Supplementary figures

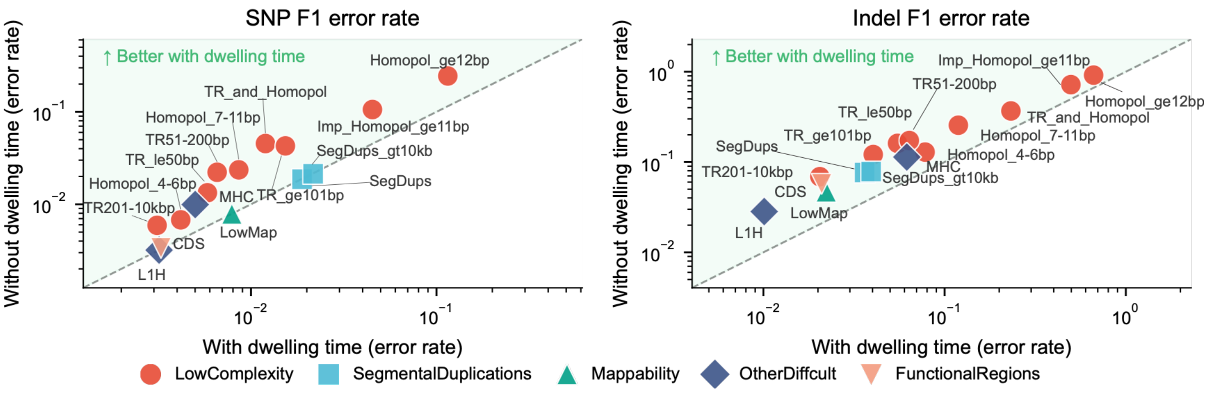

***Supplementary Figure 1 Performance in complex genomic regions (HG001).*** *F1 error rates (1 – F1) for SNPs (left) and indels (right) across 16 stratified regions on HG001 at 30× coverage. Points above diagonal indicate improvement with Clair3 v2.*

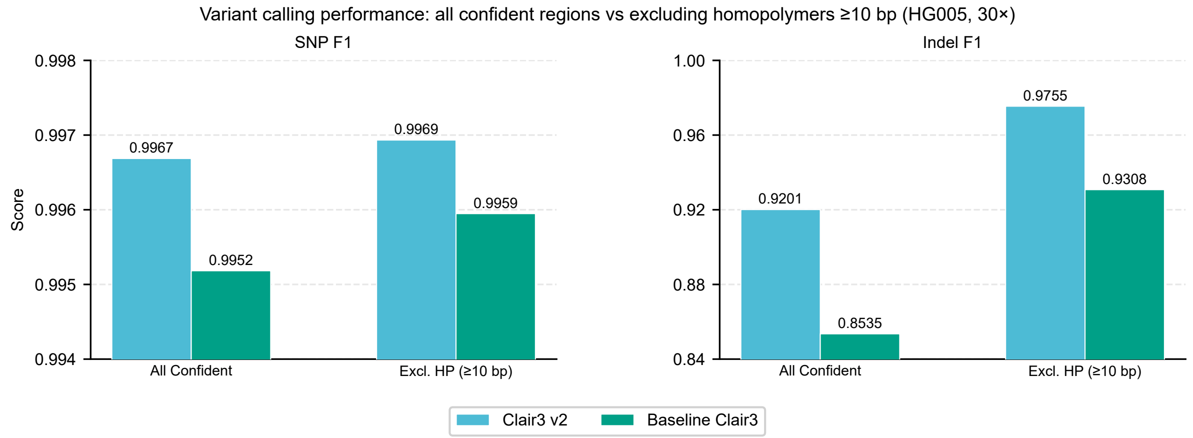

***Supplementary Figure 2 Variant calling performance with and without homopolymer regions (HG005).*** *F1 scores for SNPs (left) and indels (right) evaluated in all confident regions and after excluding homopolymers ⩾10 bp on HG005 at 30× coverage.*
